# Role of Yap1 in adult neural stem cell activation

**DOI:** 10.1101/2022.01.12.475985

**Authors:** Wenqiang Fan, Jerónimo Jurado-Arjona, Gregorio Alanis-Lobato, Sophie Péron, Christian Berger, Miguel A. Andrade-Navarro, Sven Falk, Benedikt Berninger

## Abstract

Most adult hippocampal neural stem cells (NSCs) remain quiescent with only a minor portion undergoing active proliferation and neurogenesis. The molecular mechanisms that trigger eventually the transition from quiescence to activation are still poorly understood. Here, we found the activity of the transcriptional activator Yap1 to be enriched in active NSCs. Genetic deletion of Yap1 led to a significant reduction in the relative proportion of active NSCs supporting a physiological role of Yap1 in regulating the transition from quiescence to activation. Overexpression of wild type Yap1 in adult NSCs did not induce NSC activation suggesting tight upstream control mechanisms, but overexpression of a gain-of-function mutant (Yap1-5SA) elicited cell cycle entry in NSCs and hilar astrocytes. Consistent with a role of Yap1 in NSC activation, single cell RNA sequencing revealed the partial induction of an activated NSC gene expression program. Yet, Yap1-5SA expression also induced Taz and other key components of the Yap/Taz regulon previously identified in glioblastoma stem cell-like cells. Consequently, dysregulated Yap1 activity led to repression of hippocampal neurogenesis, promoting aberrant differentiation instead.

## Introduction

During embryonic development, neural stem cells (NSCs) migrate from the dentate epithelium to settle in the dentate gyrus anlage of the hippocampus. Some of these will then eventually give rise to NSCs in the adult dentate gyrus (Berg *et al.*, 2019). A key process in the establishment of the adult NSC pool is the progressive entry of NSCs into a quiescent state in which many will persist before eventually becoming re-activated during adulthood and re-entering cell cycle (Berg *et al.*, 2019). The regulation of both transitions, i.e., the transition from actively-dividing to quiescent as well as the converse, the *de novo* activation is one of the key mysteries of adult NSC biology (Urbán *et al.*, 2019). Quiescence is defined as “reversible cell cycle arrest” and is believed to protect stem cells against damage and premature exhaustion of the stem cell pool (Urbán *et al.*, 2019). Intriguingly, maintenance of quiescence is dynamically regulated in that, during aging, the rate of activation decreases with more NSCs persisting in a quiescent state (Dulken *et al.*, 2019, Harris *et al.*, 2021, Ibrayeva *et al.*, 2021, Kalamakis *et al.*, 2019). Several signaling pathways have been implicated in the bidirectional traffic between quiescence and activation, including BMP (Mira *et al.*, 2010), Notch (Engler *et al.*, 2018, Harada *et al.*, 2021, Kawai *et al.*, 2017, Zhang *et al.*, 2019), VEGFR3 (Han *et al.*, 2015) and Wnt signaling (Lie *et al.*, 2005, Wexler *et al.*, 2009), but also include circuit-specific mechanisms (Song *et al.*, 2012). At a transcriptional level, previous work has shown that in the adult hippocampus expression of Ascl1 is a prerequisite for activation of quiescent NSCs (Andersen *et al.*, 2014), while degradation of Ascl1 protein is an important step in controlling return to quiescence (Urbán *et al.*, 2016). However, the full spectrum of transcriptional mechanisms underpinning NSC activation is still incompletely understood.

Accruing evidence points to the importance of the Hippo signaling pathway in regulating stem cell renewal and reactivation during regenerative processes in various tissues (Mo *et al.*, 2014, Moya *et al.*, 2019). The Hippo pathway is an evolutionary conserved kinase cascade which regulates the activity of its effector proteins, the transcriptional activators Yap1 (Yes-associated protein 1) and Taz, through phosphorylation, thereby causing their cytoplasmic retention and degradation (Yu *et al.*, 2013). Several mechanisms converge on this kinase cascade, including mechanical stimuli, cell polarity as well as other signaling pathways (Yu *et al.*, 2013).

Recent work has implicated the Hippo effector protein Yorkie (Yki) in the control of activation of quiescent NSCs in the brain of *Drosophila* larvae (Ding *et al.*, 2016, Gil-Ranedo *et al.*, 2019). For instance, loss of Yki has been shown to result in the failure of larval NSCs to grow in response to hormone stimulation and to re-enter cell cycle (Ding *et al.*, 2016). The Hippo pathway and the Yki orthologue Yap1 have been shown to be highly active in the developing mammalian forebrain (Cappello *et al.*, 2013, Kostic *et al.*, 2019, Lavado *et al.*, 2018, Mukhtar *et al.*, 2020), but its role in the functional state of NSCs in adult neurogenic niches has not been addressed so far.

Thus, we wondered whether Yap1 activation is an evolutionary conserved mechanism in the regulation of NSC activity. Towards this, here we have re-analysed published single cell RNA sequencing data (Hochgerner *et al.*, 2018) which revealed that Yap1 is highly enriched in activated NSCs of the adult hippocampus. This has prompted us to perform loss- and gain-of-function studies in the adult hippocampus which provide evidence for an important role of this transcriptional activator in the regulation of adult hippocampal NSC activity.

## Results

### Evidence for Yap1 activity in active neural stem cells of the adult hippocampus

To obtain first evidence for ongoing *Yap1* activity in the lineage of adult hippocampal neural stem cells (NSCs), we first re-analysed a published data set of single cell transcriptomes comprising NSCs and their early-stage progeny isolated from the dentate gyrus of the adult hippocampus (Hochgerner *et al.*, 2018). This identified a cell population enriched in a reference signature of active NSCs (Shin *et al.*, 2015), originally classified as neuronal intermediate progenitor cells (nIPCs) (Fig. EV1). We next assessed which of these cell populations, if any, exhibited expression of a reference signature of *Yap1* activity (Cordenonsi *et al.*, 2011). This revealed a specific enrichment of *Yap1* activity signature in the same cell population enriched for the active NSC signature (Fig. EV1). This analysis, thus, suggests that activation of NSC may involve Yap1-mediated transcription.

Based on this analysis, we next examined whether we could detect Yap1 protein expression within the NSC lineage (Fig. 1). We found Yap1 specifically expressed in Sox2-positive (+) cells within the subgranular zone (SGZ) (Fig. 1A) as well as hilar astrocytes (identified by GFP expression driven from human glial fibrillary protein (hGFAP) gene regulatory elements and hilar localisation) (Fig. 1B), but virtually absent in doublecortin (DCX)+ young neurons (Fig. 1C) or Olig2+ cells (i.e., comprising cells of oligodendroglial lineage) (Fig. 1D). While in hilar astrocytes Yap1 immunoreactivity appeared to be largely cytoplasmic, within the SGZ, some of the Yap1 immunoreactivity co-localized with Sox2 expression, demarcating the nuclei of these Sox2+ cells (Fig. 1A, inset). This data suggests that Sox2+ SGZ cells (i.e., NSCs and intermediate progenitor cells) are enriched in Yap1 expression, and that Yap1 may have undergone nuclear translocation in a subpopulation of these cells.

**Figure 1.**
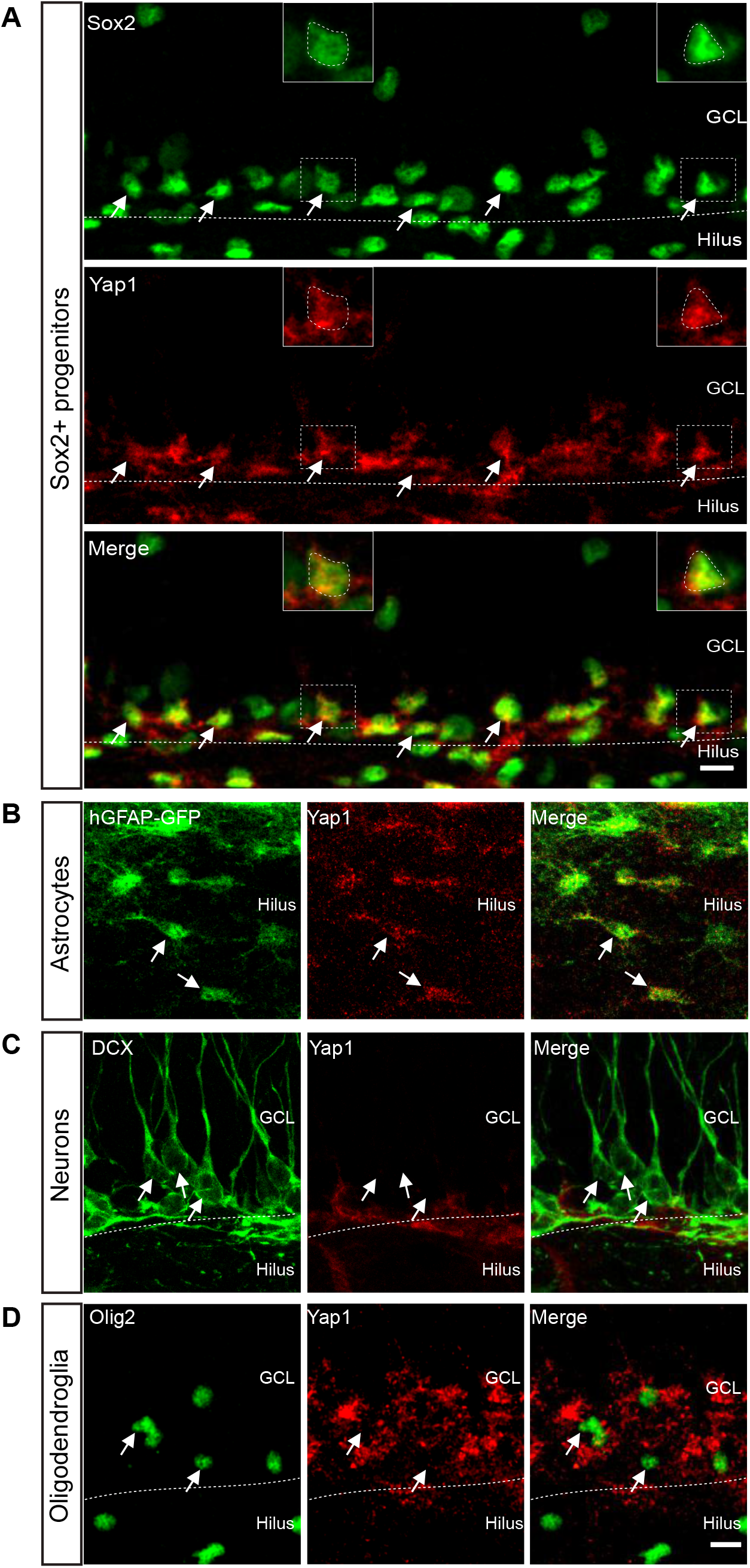
Expression pattern of Yap1 in the DG of the adult hippocampus. A. Representative images of immunostainings of adult mouse brain coronal sections showing that Yap1 (red) is expressed in Sox2-positive (green) neural progenitors of SGZ in the DG. Arrows indicate double-positive cells. B. Immunofluorescence for GFP and Yap1 in adult brain coronal sections from hGFAP-GFP transgenic mice. Arrows indicate GFP-positive astrocytes in the hilus expressing Yap1. C-D. Immunofluorescence for DCX, Olig2 and Yap1 in adult brain coronal sections. Arrows indicate DCX-positive immature neurons and Olig2-positive oligodendroglial cells. Data information: GCL, granule cell layer. Scale bars: 10 µm.

In order to assess whether NSC activation is associated with nuclear enrichment of Yap1, we took advantage of primary adult hippocampal NSC cultures (Babu *et al.*, 2011) (Fig. EV2A), which have been successfully used to study the transition between NSC states of activation and quiescence (Martynoga *et al.*, 2013) (Fig. EV2B-D). Immunofluorescence analysis for Yap1 location revealed a clear enrichment in nuclear Yap1 in active versus quiescent NSCs (Fig. EV2E and F). This data supports the notion that NSC activation is accompanied by transfer of Yap1 from the cytoplasm to the nucleus.

### Long-term *Yap1* loss-of-function compromises NSC activation

To study the role of *Yap1* in regulating the NSC transition from quiescence to activation, we next analyzed the impact of *Yap1* loss-of-function on the ratio of quiescent and active NSCs. Towards this, we deleted *Yap1* specifically in NSC (and astrocytes) and their lineage by crossing *Yap1* conditional knock-out mice (Yap1fl/fl) (Zhang *et al.*, 2010) with Glast-CreERT2 and CAG-CAT-EGFP mice (Mori *et al.*, 2006, Nakamura *et al.*, 2006) followed by treatment with tamoxifen (Fig. 2A). Immunofluorescence for Yap1 revealed reduction of expression by day 7 following induction of recombination, indicating successful *Yap1* deletion (Fig. 2B and C). To assess the proportion of actively dividing cells among NSCs 7 days after induction of recombination, we first determined the number of recombination-reporter-positive (GFP+) cells among radial glia-like cells (RGLs, i.e., the major NSC population, identified by their localization in the SGZ, GFAP immunoreactivity, and the presence of a radially-oriented process) and then quantified those RGLs that were engaged in cell cycle (minichromosome maintenance complex component-2, Mcm2-positive cells) (Fig. 2D). Quantitative, double-blind analysis revealed that the proportion of active versus quiescent (Mcm2+ versus Mcm2-) RGLs had not changed at day 7 following deletion of *Yap1* as compared to controls (Fig. 2E and F). Likewise, when we performed the same analysis 30 days after inducing the deletion of *Yap1*, we did not observe significant changes of active versus quiescent RGLs (Fig. EV3). By contrast, at day 60 following deletion of *Yap1*, we noted a significant drop in the number of active RGLs as compared to controls (Fig. 2G-I). This set of experiments indicates that loss of *Yap1* does not immediately impact on the levels of NSC transition from quiescence to activation, but that long-term *Yap1* loss-of-function results in decreased NSC activation, evidencing an important physiological role of *Yap1* in this process.

**Figure 2.**
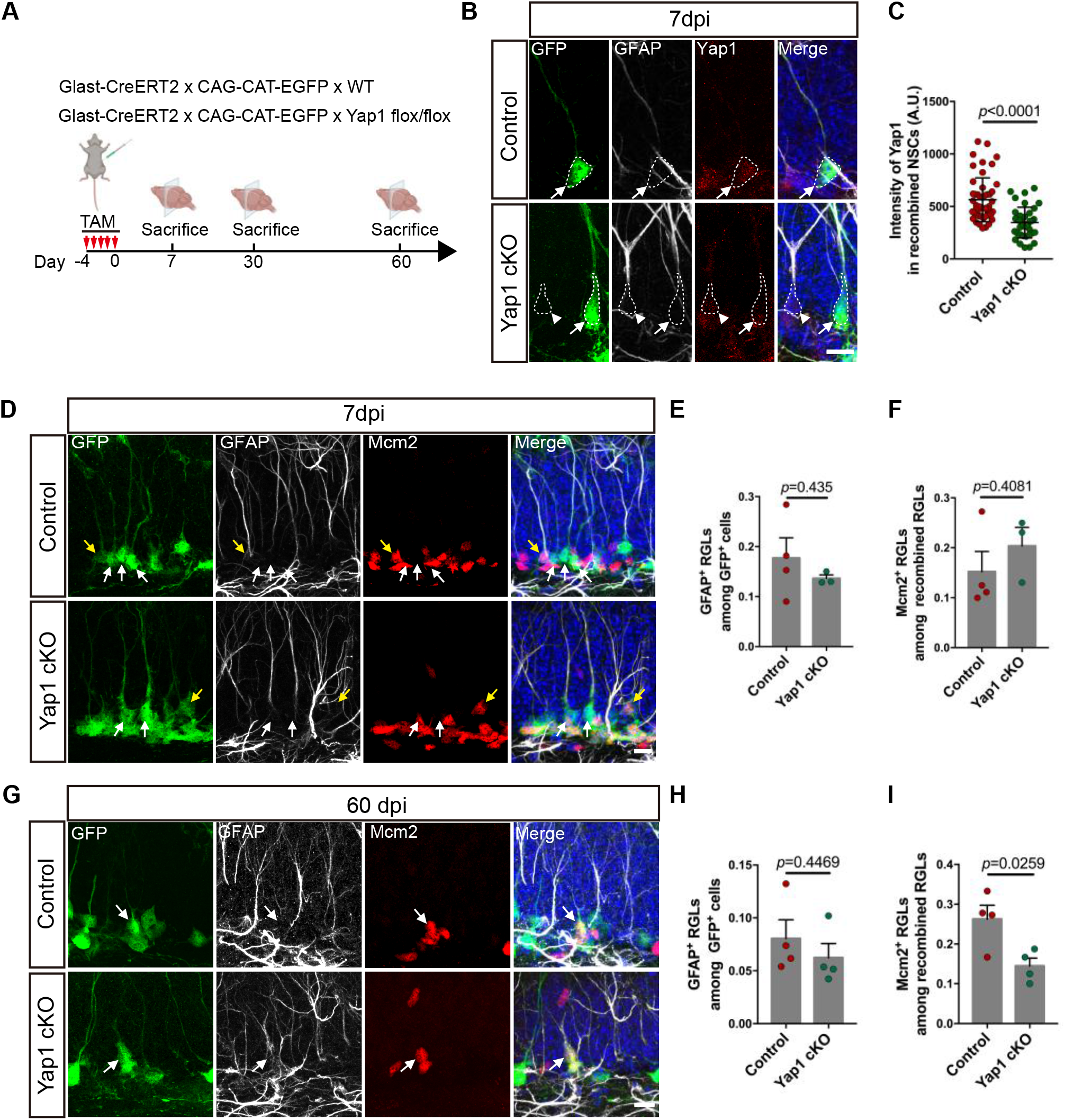
Yap1 regulates the balance between quiescent and active NSCs. A. Schematic diagram of the experimental design. Adult (8 week-old) Yap1 cKO mice or Control mice were injected with Tamoxifen for 5 consecutive days and sacrificed 7, 30, or 60 days later for immunohistological analysis. B. Immunofluorescence for GFP (recombination reporter), GFAP and Yap1 in control and Yap1 cKO RGLs at 7 days after tamoxifen administration. Arrowhead indicates non-recombined RGL and arrow indicates recombined RGL. C. Quantification of Yap1 intensity in (B). Yap1 level is significant decreased in Yap1 cKO RGLs at 7dpi. n= 50 cells from 3 mice in control group, n= 37 cells from 3 mice in Yap1 cKO group. D. Immunofluorescence for GFP, GFAP and Mcm2 in control and Yap1 cKO RGLs at 7 days after tamoxifen administration. Yellow arrow indicates proliferating adult RGL, white arrows indicate quiescent adult RGLs. E-F. Quantification of the data in (D). Loss of Yap1 does not induce any changes neither in the number of proliferative RGLs nor the overall number of RGLs. n=4 mice for control group, n=3 mice for Yap1 cKO group. G. Immunofluorescence for GFP, GFAP and Mcm2 in control and Yap1 cKO RGLs at 60 days after tamoxifen administration. Arrows indicate Mcm2-positive, proliferating RGLs. H-I. Quantification of the data in (G). Loss of Yap1 results in the decrease of proliferative RGLs, while the overall number of RGLs does not change. n=4 mice in control, n=4 mice in Yap1 cKO. Data information: Data are represented as mean ± SEM. Unpaired Student’s t test. Scale bars: 10 µm. dpi=days post injection.

### Active Yap1 promotes cell cycle entry of quiescent hippocampal NSCs

We next investigated the effect of *Yap1* gain-of-function on the activation state of adult hippocampal NSCs. Towards this, we targeted adult NSCs (as well as hilar astrocytes) using lentiviruses driving transgene expression under the control of hGFAP regulatory elements (LV-hGFAP-IRES-EGFP, LV-hGFAP-Yap1WT-IRES-EGFP, LV-hGFAP-Yap1-5SA-IRES-EGFP) (Fig. EV4). Interestingly, 7 days following lentivirus-mediated expression of the wild type form of *Yap1* (Yap1 WT) in adult hippocampal SGZ, no significant change in the number of proliferative Sox2+ progenitors (i.e. lentivirus-transduced Sox2+ cells in the SGZ) was observed (Fig. 3A, B and C). In contrast, lentivirus-mediated expression of a mutant form of *Yap1*, encoding a protein in which the serines of the five HXRXXS Lats motifs had been replaced by alanine (Yap1-5SA), rendering the protein insensitive to phosphorylation-dependent inhibition by Lats and thereby constitutively active (Zhao *et al.*, 2007), caused a dramatic rise in the number of lentivirus-transduced Ki67+Sox2+ cells (Fig. 3B and C). Finally, we also observed an overall increase in the density of Sox2+ cells in the SGZ (Fig 3B and D). This data shows that active Yap1 is sufficient to drive quiescent NSCs towards cell cycle entry. It also supports the notion that Yap1 activity is under strict control as forced expression of Yap1 WT fails to induce NSC activation, and only a regulation-deficient form of Yap1 overcomes this control. In line with the Yap1-mediated activation of hippocampal NSC *in vivo*, we found that lentivirus-mediated expression of Yap1-5SA promoted dramatic cell cycle entry in primary cultures of hippocampal NSCs kept in quiescence conditions as assessed by phosphohistone-H3 staining as well as EdU incorporation (Fig. EV5 A-D).

**Figure 3.**
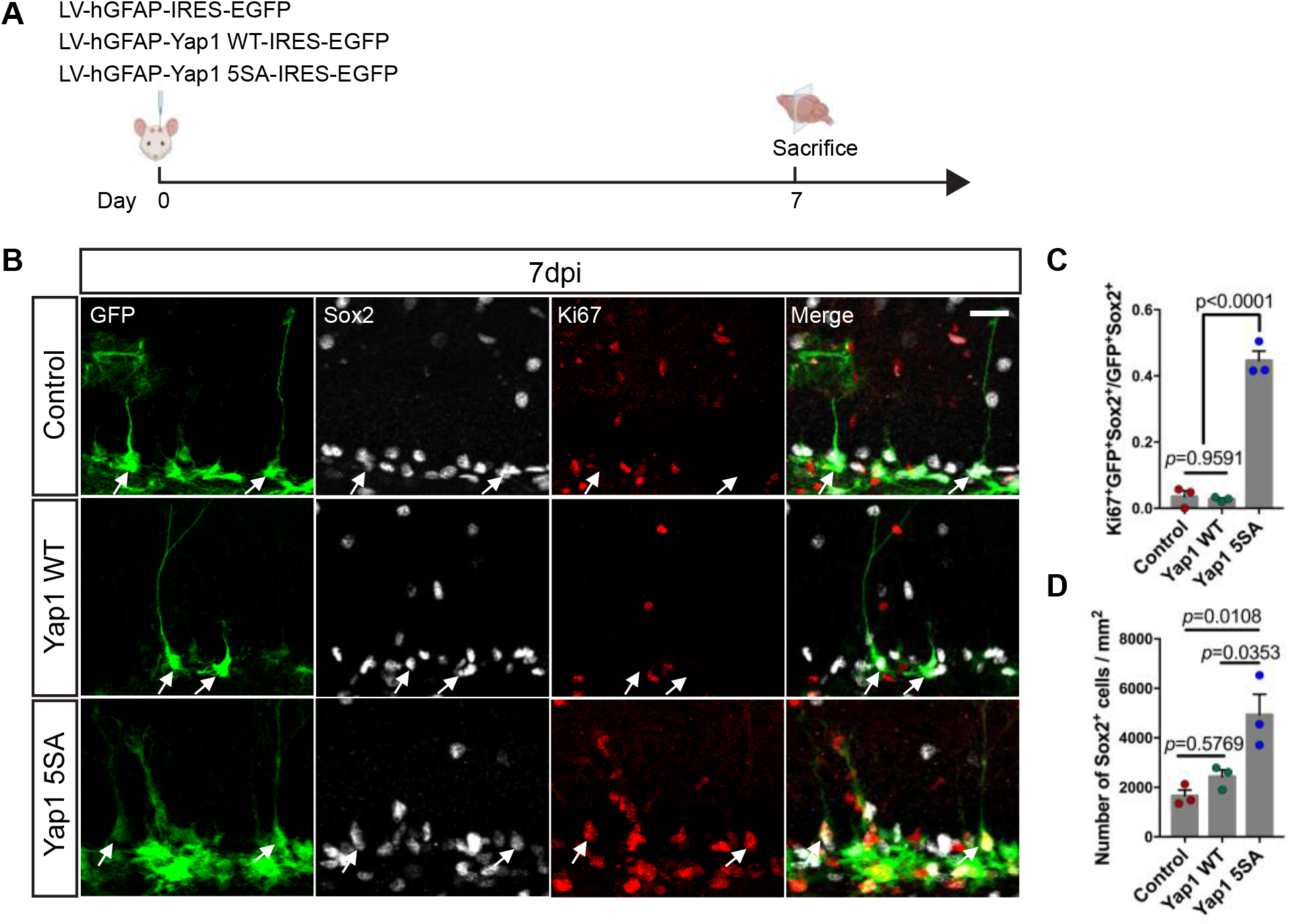
Yap1 gain-of-function induces adult hippocampal NSC proliferation *in vivo*. A. Schematic diagram of experimental design. P60 mice were analyzed at 7 days after lentivirus injection of control, wild type Yap1 and Yap1-5SA. B. Immunofluorescence for GFP, Sox2 (to identify NSCs, arrows) and Ki67 (to identify proliferative cells) in control conditions or overexpression of wildtype Yap1 or Yap1-5SA 7days after lentivirus injection. C-D. Quantification of proliferative NSCs. Yap1-5SA induces the proliferation of adult NSCs and increases the number of Sox2-positive cells. n=3 mice for each group. Data information: Data are represented as mean ± SEM. Unpaired Student’s t test. Scale bars: 50 µm.

Finally, in these injections some astrocytes in the hilus of the dentate gyrus were targeted by the lentiviruses encoding GFP only or Yap1-5SA and GFP. While we did not observe any Ki67 positive astrocytes following control virus injection, many astrocytes were Ki67 positive following expression of Yap1-5SA. This indicates that Yap1-5SA also stimulates cell cycle re-entry in postmitotic astrocytes (Fig. EV5E and F).

### Constitutively active Yap1 induces an aberrant active NSC-like signature in adult hippocampal NSCs and/or astrocytes

Given our observation that forced expression of Yap1-5SA appears to activate quiescent NSCs, we finally addressed how this molecular perturbation alters gene expression using single cell RNA-sequencing. Yap1-5SA was targeted to adult hippocampal NSCs and hilar astrocytes by injection of lentiviruses encoding Yap1-5SA and EGFP or EGFP alone for control, under hGFAP regulatory elements, into the SGZ of the adult dentate gyrus. We performed fluorescence-activated cell sorting (FACS) to isolate EGFP-positive cells at day 3 and 7 following lentivirus transduction (Fig. EV6). We selected isolation at day 3 as this was the earliest time point when we could detect reliable EGFP expression, while day 7 corresponds to the time point when we had previously observed massive activation of quiescent NSCs (Fig. 3). Single cell transcriptomes were measured following library preparation using Smart-seq2 methodology (Fig. 4A) (Picelli *et al.*, 2014). This yielded 121 (day-3) and 258 (day-7) control and 273 (day-3) and 320 (day-7) Yap1-5SA-expressing single cell transcriptomes. Using Scanpy (Wolf *et al.*, 2018) we identified 13 distinct clusters (Fig. EV7A) and employed force-directed graph embedding for visualization. Clusters expressing astrocyte, quiescent NSC, neuron, oligodendrocyte precursor cell (OPC), oligodendrocyte, and microglia markers consisted exclusively of control cells (Fig. EV7). Conspicuously, none of these clusters contained cells expressing Yap1-5SA, suggesting that this perturbation exerts a drastic effect on the targeted cells forcing them to alter their cell state within a 3-day time window. Among the remaining clusters, several contained both control and Yap1-5SA-expressing cells, while a final set of clusters consisted of Yap1-5SA cells only. Consistent with Yap1-5SA being a transcriptional activator (Zhao *et al.*, 2007), clusters containing Yap1-5SA-expressing cells exhibited increased number of transcripts compared to control cell clusters (Fig. EV7B). To further reveal the identity of Yap1-5SA-expressing cells and their lineage relationship to quiescent NSC and astrocytes, we performed re-clustering after removal of microglia, OPCs, oligodendrocytes, as well as a cluster enriched in cell-death related genes (Mdm2, Trp53), while the cluster identified with neurons was retained as main natural progeny of the adult hippocampal NSC lineage (Fig. 4B-F). This re-analysis yielded 12 clusters comprising astrocytes, quiescent NSCs, neurons, none of which contained Yap1-5SA cells, and 9 clusters entirely consisting of Yap1-5SA-expressing cells (Fig. 4B and E, Fig. EV8). To obtain insights into the lineage dynamics between the different clusters, we employed RNA velocity with scvelo (Bergen *et al.*, 2020, La Manno *et al.*, 2018), supporting the notion that the overall lineage progression vector originated from the cluster comprising the NSC population. (Fig. 4B and E). Indeed, pseudotime analysis identified NSCs as source for all other cells, with neurons representing the most distant differentiated fate (Fig. 4C and E), consistent with the natural lineage progression of adult hippocampal neurogenesis. As expected, control-transduced cells showed low levels of gene expression related to cell cycle (*cell-cycle signature* (Hao *et al.*, 2021), Fig. 4D) or Yap1 activity (*Yap1 signature* (Cordenonsi *et al.*, 2011), Fig. 4D). Moreover, only few cells were enriched in gene expression related to an activated NSC state (*activated NSC signature* (Shin *et al.*, 2015), consistent with the fact that most adult hippocampal NSCs are quiescent (*quiescent NSC signature* (Shin *et al.*, 2015), Fig. 4D and EV8B). In contrast, following Yap1 gain-of-function, we observed an enrichment in Yap1 signature across all Yap1-5SA-expressing clusters (Fig. 4D). Yap1-5SA-expressing clusters transcriptionally closer to control-transduced NSCs exhibited elevated cell-cycle signature (Fig. 4D). Accordingly, these clusters (2,4-7) down-regulated genes enriched in quiescent NSCs (Fig. 4D), while up-regulating an activated NSC signature (Fig. 4D, Fig. EV8B). This data is thus consistent with the notion that Yap1 gain-of-function can elicit activation of quiescent NSCs and cell-cycle entry, as observed in our *in vivo* and *in vitro* experiments (Fig. 3 and EV2). One intriguing observation was the absence of significant Ascl1 expression in Yap1-5SA-expressing cells, in sharp contrast to control NSCs, consistent with the fact that the former cells failed to follow a trajectory leading to adult neurogenesis. While Yap1-5SA-expressing did not contribute to neuronal fate (Fig. EV8B), we observed at least two additional RNA velocity vector-endpoints (clusters 9 and 11) comprising exclusively Yap1-5SA-expressing cells and characterised by reduced cell-cycle and activated NSC signatures (Fig. 4C and D, Fig. EV8B). Intriguingly, these clusters were typified by high levels of Scg2 (cluster 11) and Pmp22 (cluster 9) expression, respectively (Fig. 4E). However, these cells also expressed an arsenal of non-neural-specific genes, suggestive of aberrant differentiation (Fig. EV8A). Also, while less segregated in pseudotime, Yap1-5SA-expressing cluster 1 also showed reduced cell-cycle signature and exhibited a prominent expression of mesenchymal cell type-related genes (e.g., Col1a1, Fig. 4D, E and EV8A). Given the strong implication of Yap1 in brain tumour initiation, we finally scored control and Yap1-5SA-expressing cells for their expression of glioblastoma stem cell-related genes (*glioblastoma stem cell-like signature* (Castellan *et al.*, 2021), such as the Yap1 paralogue transcriptional regulator Taz as well as some of their primary downstream transcriptional regulators (Fig. 4F). While Yap1-5SA-expressing cells were overall enriched in glioblastoma stem cell-like signature across clusters, cluster 9 and 11 showed lower enrichment consistent with their partial, albeit aberrant differentiation. Finally, we noted that most Yap1-5SA-expressing clusters comprised cells isolated on days 3 or 7, but enrichment of day-7 cells in clusters 2 and 1 (Fig. EV8B). This may suggest that progression towards a more mesenchymal phenotype may involve prolonged Yap1 activity. In sum, our scRNA-seq data indicate that Yap1 gain-of-function in adult hippocampal NSC and astrocytes can induce molecular hallmarks of an activated NSC. However, prolonged Yap1 activity disrupts the physiological molecular trajectory towards neurogenesis and instead induces a gene expression signature akin to glioblastoma stem cells and can lead to aberrant differentiation.

**Figure 4.**
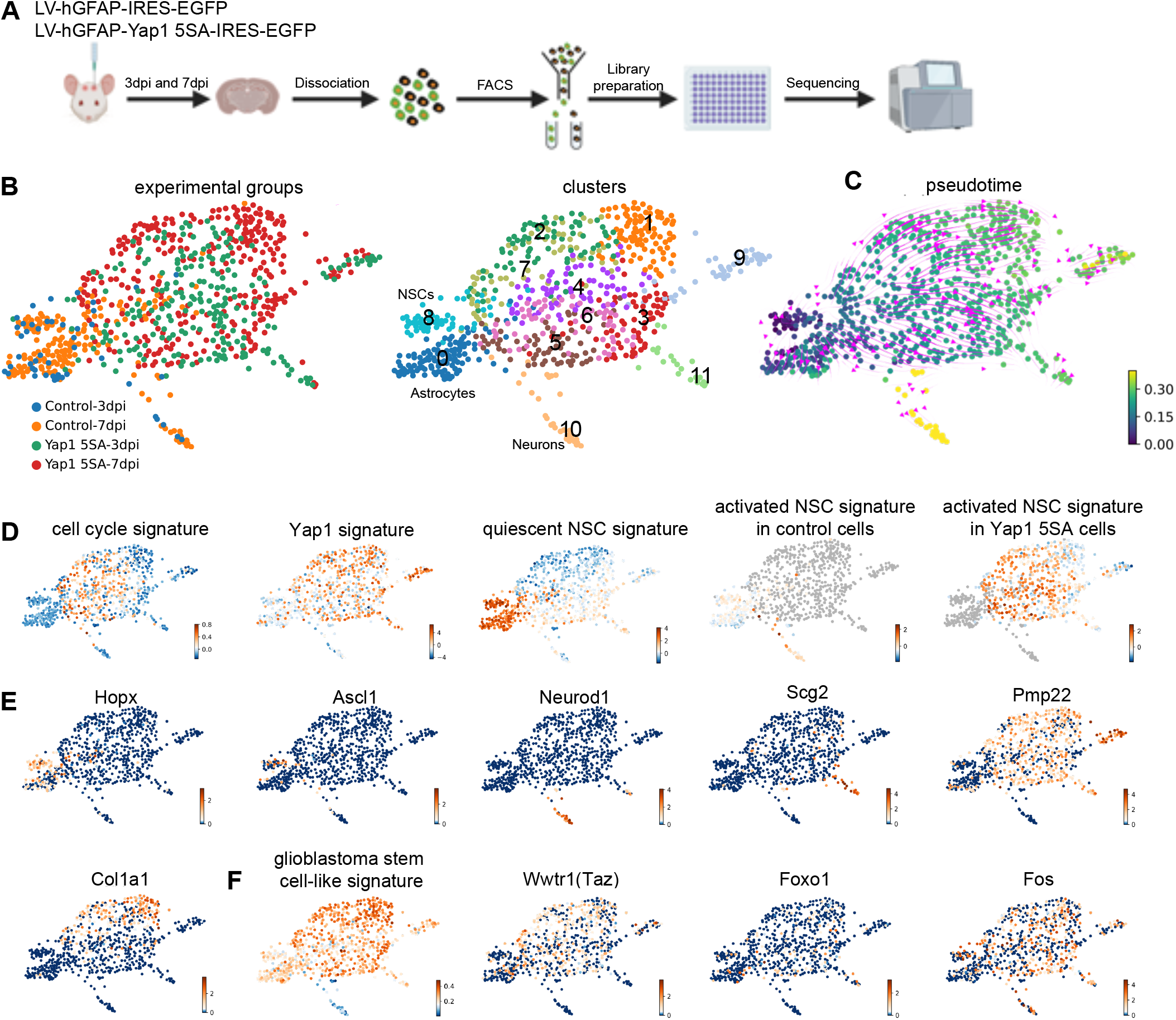
Overexpression of active Yap1 induces an aberrant active NSC-like signature. A. Schematic diagram of experimental design of single cell RNA sequencing at 3 and 7 days after viral injection. B. Clustering after force-directed graph embedding and removal of cells with microglia, OPCs, oligodendrocyte or cell death signatures. C. Diffusion pseudotime and RNA velocity analysis indicating cell fate trajectories. D. Force-directed graph plots showing signatures of cell cycle-related, Yap1 activity-related, quiescent NSC-related and activated NSC-related gene expression. E. Force-directed graph showing expression of Hopx, Ascl1, Neurod1, Scg2 and Pmp22 and Col1a1. F. Force-directed graph displaying glioblastoma stem cell-like signature and expression of Yap/Taz hub genes Wwtr1 (Taz), Foxo1, and Fos.

## Discussion

In this study, we examined the hypothesis that the transcriptional co-activator Yap1 plays a role in the transition of adult hippocampal NSCs from quiescence to activation. Consistent with this hypothesis, we find that Yap1 expression is enriched in activated NSCs and YAP1 gain-of-function can trigger cell cycle entry in quiescent NSCs. Conversely, Yap1 gene deletion in adult hippocampal NSCs results in a reduction in the ratio of activated vs. quiescent NSCs. Finally, prolonged Yap1 activity in adult hippocampal NSCs disrupts physiological neurogenesis promoting aberrant cell differentiation and partial acquisition of a glioblastoma stem cell-like signature.

Our study set out from the re-analysis of published scRNA-seq data (Hochgerner *et al.*, 2018) showing that Yap1 activity is highly enriched in activated NSCs, suggesting that Yap1 may play an important role as regulator of the transition of adult hippocampal NSCs from a quiescent to an activated state. A key step in the regulation of Yap1 activity consists in its translocation from the cytoplasm to the nucleus. Indeed, we did not only find that Yap1 is selectively expressed in adult hippocampal NSCs *in vivo*, but a hippocampal NSC culture allowed us to reveal that activation of quiescent NSCs is accompanied by a marked nuclear translocation of Yap1 protein. Intriguingly, *in vivo* overexpression of wildtype Yap1 did not elicit overt NSC activation. This data indicates that in our *in vivo* experiments, regulatory mechanisms upstream of Yap1 were not overwhelmed by the elevated Yap1 protein levels, suggesting tight control over wild type Yap1 activity in adult hippocampal NSCs under normal conditions. Interestingly, when quiescent NSCs in culture are experimentally activated by removal of BMP4 and treatment with EGF/FGF2 (Martynoga *et al.*, 2013), NSCs are relieved from this tight control, as illustrated by the drastic increase in nuclear Yap1 protein. In contrast to wild type Yap1, overexpression of a Yap1 mutant defective for phosphorylation-mediated inhibition (Yap1-5SA) (Zhao *et al.*, 2007), induced cell cycle entry in adult hippocampal NSCs. This clearly shows the potency of Yap1 in promoting NSC activation once relieved from inhibitory control. Likewise, we found that hilar astrocytes commence to proliferate in response to Yap1-5SA overexpression.

In the present study, we did not address the upstream mechanisms responsible for the effective control of Yap1 activity in NSCs or astrocytes. However, Müller glia in the mouse retina have been shown to exit quiescence and commence proliferation upon deletion of Lats1 and 2, i.e., key components of the canonical Hippo pathway, indicating that Hippo signaling exerts inhibitory control under physiological conditions (Hamon *et al.*, 2019, Rueda *et al.*, 2019). While it is likely that similar mechanisms also operate in the adult NSC niche, it remains to be shown what molecular signals converge onto the Hippo pathway to maintain Yap1 activity at bay.

Given the apparent tight control of Yap1 in adult hippocampal NSCs, we were somewhat surprised to see the initially only very subtle effect of Yap1 deletion (i.e., at seven or 30 days after deletion). A significant change in the ratio of activated vs. quiescent NSCs was only detected after 60 days post deletion. One reason for this delayed response may consist in the temporary compensation by the Yap1 paralogue Taz (Plouffe *et al.*, 2018). A second reason may be related to the fact that even with early aging, the frequency of quiescent NSCs to become activated decreases significantly (Harris *et al.*, 2021, Ibrayeva *et al.*, 2021, Kalamakis *et al.*, 2019), possibly due to downregulation of activation-promoting and/or upregulation of quiescence-retaining mechanisms (Harris *et al.*, 2021). Such changes could render the activation of quiescent adult hippocampal NSCs increasingly dependent on Yap1 activity and thereby enhance the consequences of Yap1 deletion compared to earlier stages. In any case, the physiological requirement of Yap1 in adult NSC activation observed in our study is highly reminiscent of the impaired reactivation of Müller glia in the injured adult mouse retina following Yap1 gene deletion (Hamon *et al.*, 2019, Rueda *et al.*, 2019). Also, given its proliferation-promoting effect on hilar astrocytes, it will be interesting to decipher the physiological contribution of Yap1 in the induction of proliferation of parenchymal astrocytes during reactive gliosis.

Compared to its role in regulating adult NSC activity, Yap1 activity appears to play an even more prominent role during embryonic neurogenesis. Using Yap1 loss-of-function approaches, Kostic et al. observed a significant reduction in the proliferation of basal progenitors in ferret and human cortex. Intriguingly, there seems to be an increase in nuclear Yap1 levels alongside with an increase in outer radial glia (ORG) from lissencephalic (mouse) to gyrencephalic (ferret and human) brains, implicating differential regulation of Yap1 activity in brain evolution. Recent data suggest a close relationship between ORG and adult hippocampal NSCs (Berg *et al.*, 2019). Future studies may address whether differential Yap1 activity contributes to differences in initial pool size, maintenance, activation rates, and age-dependent exhaustion of adult hippocampal NSCs across mammalian species.

Consistent with a role of Yap1 in inducing NSC activation, scRNAseq from Yap1-5SA-expressing NSCs and astrocytes revealed a reduction in the expression of genes normally transcribed in quiescent NSCs and a concomitant increase in genes related to cell cycle and NSC activation. However, we noted that both at day 3 and day 7, transcriptomes of Yap1-5SA-expressing cells deviated substantially from the trajectory leading to neurons, indicative of a more profound and most likely aberrant reprogramming of gene expression. One particularly intriguing observation was the conspicuous absence of Ascl1 expression in Yap1-5SA-expresing cells. Onset of Ascl1 expression is a hallmark of adult hippocampal NSC activation and its genetic deletion results in NSC activation failure (Andersen *et al.*, 2014, Urbán *et al.*, 2016). While our data indicate that Ascl1 expression may not be an absolute prerequisite for entering cell cycle, it is likely critical for endowing proliferative NSCs with a neurogenic program. It would be interesting to learn in the future why Yap1 gain-of-function results in suppression of Ascl1 induction. Intriguingly, Yap1-5SA-expressing cells may engage with various alternative differentiation programs as indicated by pseudotime analysis incorporating RNA splicing information. Partial differentiation of Yap1-5SA-expressing cells is further supported by the decrease in the expression of cell-cycle-relevant genes despite an elevated Yap1 activity signature. One peculiar observation is that Yap1-5SA-expressing cells can adopt distinct differentiated states, highlighted by the differential expression of Scg2 and Pmp22 in subpopulations of Yap1-5SA-expressing cells. Also, cluster 1 appeared to be another (less clearly segregated) endpoint in pseudotime, enriched in collagen gene expression, suggestive of acquisition of a more mesenchymal phenotype. However, none of these states could be directly identified with existing cell types. Whatever, the differentiation programs induced in Yap1-5SA-expressing cells, canonical lineage progression towards neurogenesis was severely disrupted. This suggests that sustained Yap1 activity occludes a canonical neurogenic program which is in line with the fact that Yap1 activity signature as well as Yap1 protein expression was downregulated in differentiating immature neurons. Moreover, prolonged Yap1 gain-of-function in astrocytes seems to drive these cells away from their native lineage. While entirely conceivable, our scRNAseq analysis did not allow us to distinguish whether specific Yap1 gain-of-function cell clusters were preferentially or selectively generated from adult hippocampal NSCs or hilar astrocytes.

The highly aberrant transcriptional programs elicited by Yap1 gain-of-function may reflect processes that occur during glioblastoma formation. Work by Castellan et al. (2021) recently showed that Yap1 and Taz are master regulators of stemness in glioblastoma stem-like cells (GSCs) and their gain-of-function represent a roadblock to GSC differentiation. Indeed, several of the genes identified by Castellan et al. as components of the Yap/Taz regulon in GSCs were also induced following Yap1-5SA expression in hippocampal NSCs and/or astrocytes in our study (Fig. 4F). This included the Yap1 paralogue Taz (Wwtr1) itself as well as FoxO1and Fos. Consistent with these findings, Yap1-5SA-expressing cells exhibited significant levels of expression of glioblastoma stem cell-related genes (Fig. 4F). Future studies may reveal whether dysregulation of normally tightly-regulated Yap1 activity induces the conversion of adult NSCs or postmitotic astrocytes into deadly GSCs.

## Supplementary figure legends

**Figure EV1.**
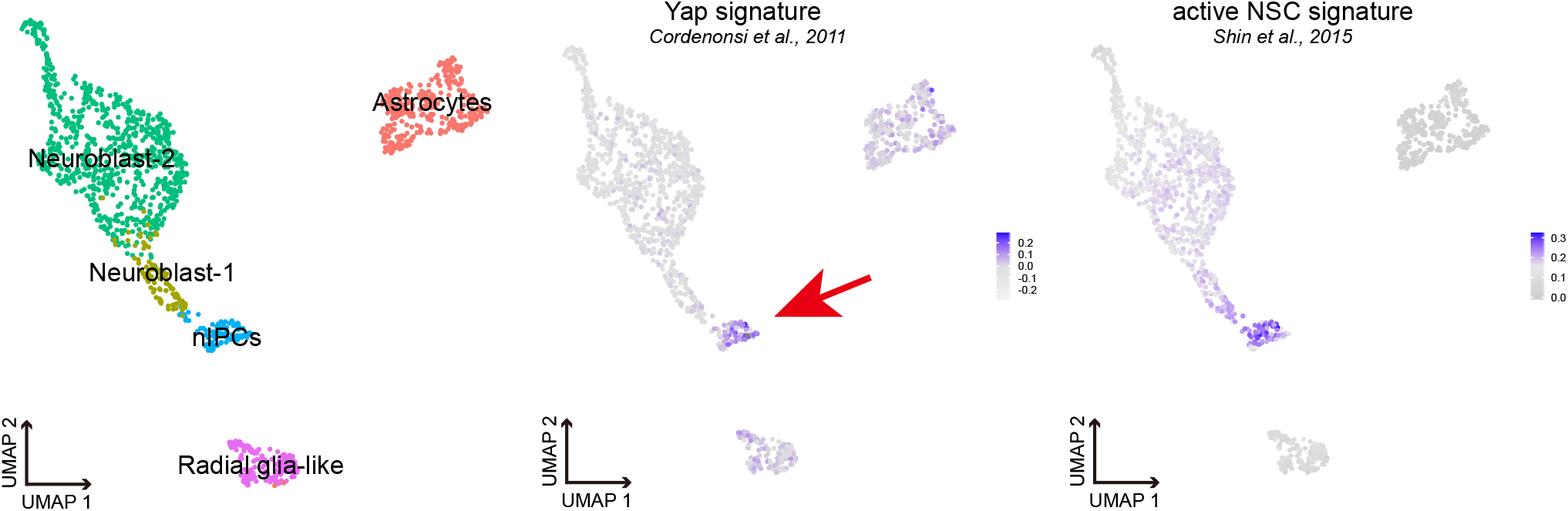
Yap1 signature is enriched in the proliferative neural progenitors of the mouse hippocampus. Uniform Manifold Approximation and Projection (UMAP) of single cell transcriptome from published data set comprising NSCs and their early progeny isolated from the dentate gyrus of the adult mouse hippocampus. Yap1 signature is enriched in the same cell population as the active NSC signature. Arrow indicates Yap1 signature in the nIPCs (neural intermediate progenitor cells).

**Figure EV2.**
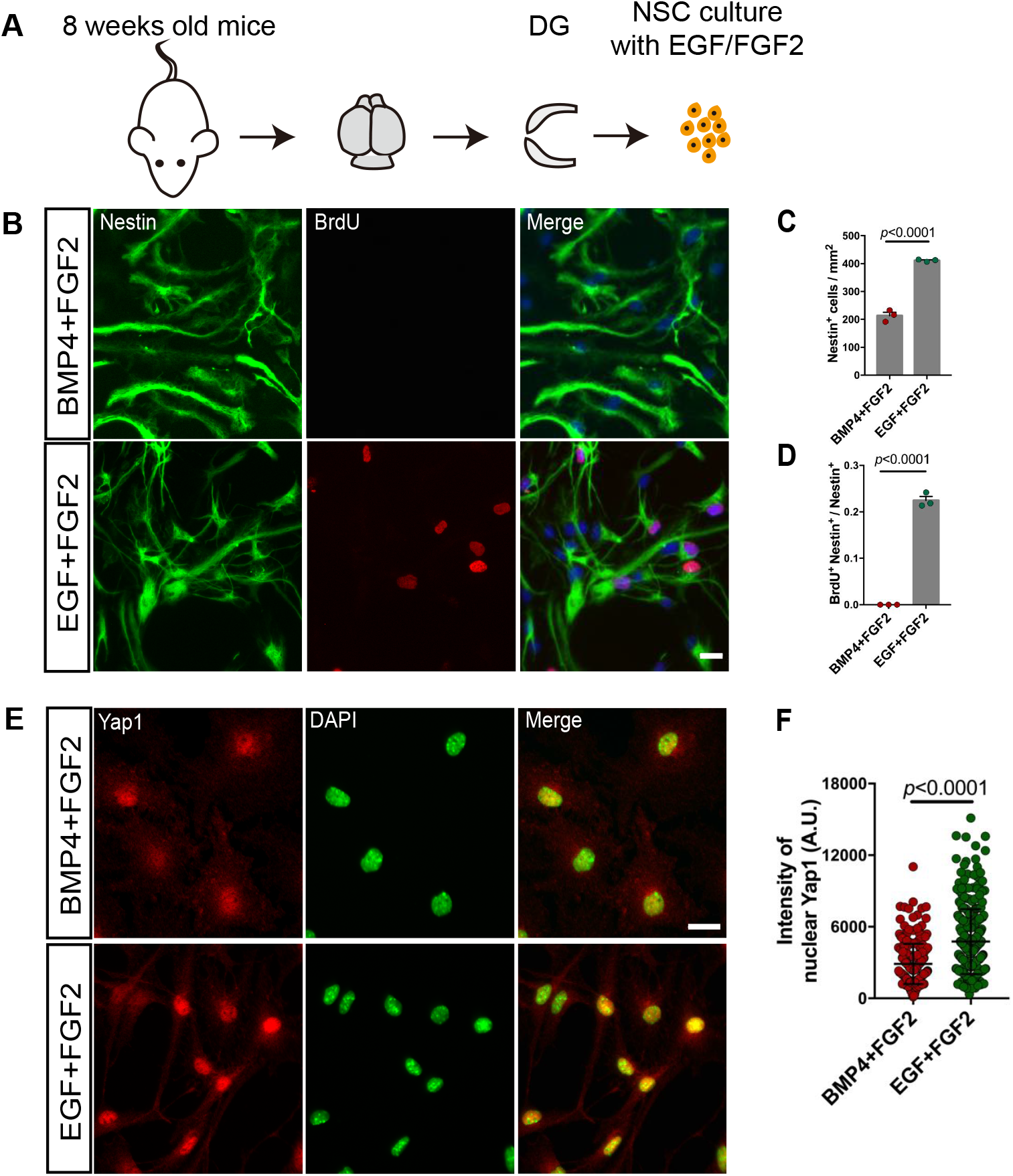
Nuclear Yap1 level is increased upon adult NSCs activation. A. Schematic diagram of adult hippocampus derived NSCs culture. B. Cultured adult NSCs were treated with EGF+FGF2 (proliferative condition) or BMP4+FGF2 (quiescent condition). Immunofluorescence for Nestin and BrdU indicates that BMP4+FGF2 treatment efficiently blocks the cell division of adult NSCs, as evidenced by the fact that they did not incorporate the BrdU administered during the last 6 hours prior to fixation. C-D. Quantification of the data in (B). No BrdU positive cells were found in quiescent condition and resulted in a reduction of NSCs density. n=3 independent experiments. E. Immunofluorescence for Yap1 and DAPI in quiescent and proliferative NSCs. F. Quantification of the nuclear Yap1 intensity in (E). Nuclear Yap1 level increases upon activation of quiescent NSCs. n=300 cells in proliferative condition, n=249 cells in quiescent condition. Data information: Data are represented as mean ± SEM. Unpaired Student’s t test. Scale bars: 20 µm.

**Figure EV3.**
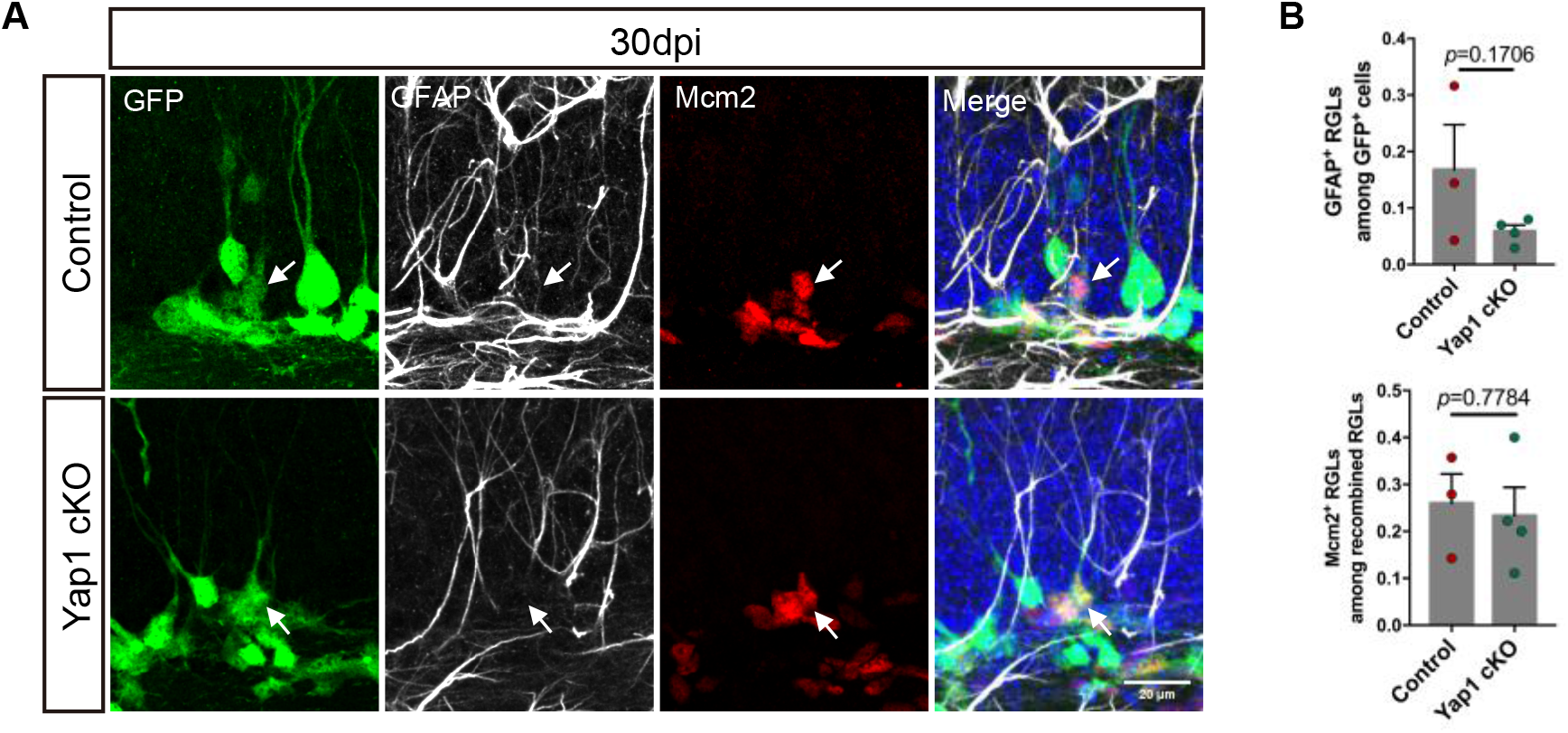
Conditional knockout Yap1 in adult NSCs at 30dpi. A. Immunofluorescence for GFP (recombination reporter), GFAP and Mcm2 in control and Yap1 cKO RGLs at 30 days after tamoxifen administration. Arrows indicate Mcm2-positive, proliferating RGLs. B. Quantification of the data in (A). Loss of Yap1 does not induce significant changes in the number of proliferative RGLs or the overall number of RGLs. n=3 mice for control group, n=4 mice for Yap1 cKO group. Data information: Data are represented as mean ± SEM. Unpaired Student’s t test. Scale bars: 20 µm.

**Figure EV4.**
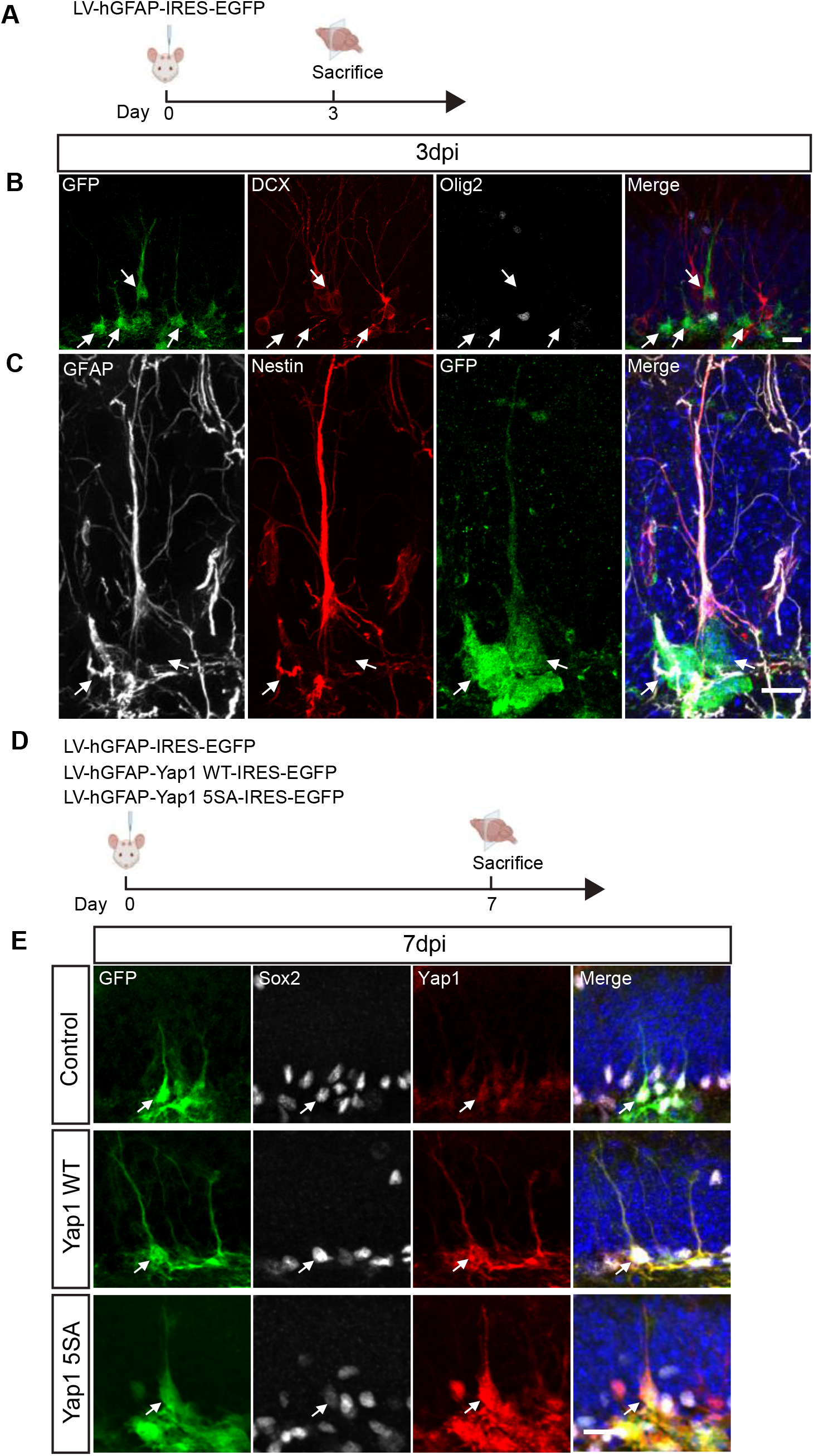
Targeting of adult NSCs *in vivo* with lentivirus under the control of hGFAP gene regulatory sequence. A. Schematic diagram of experimental design. P60 mice were analyzed at 3 days after control lentivirus injection. B. Immunofluorescence for GFP, DCX and Olig2 in coronal brain sections. Arrows indicate adult RGLs in the DG. C. Immunofluorescence for GFP, Nestin and GFAP in coronal brain sections. Arrows indicate GFAP, Nestin and GFP positive adult RGLs in the DG. D. Schematic diagram of experimental design. P60 mice were analyzed at 7 days after lentivirus injection of control, wild type Yap1 and Yap1-5SA. E. Immunofluorescence for GFP, Sox2 and Yap1 in coronal brain sections. Arrows indicate the expression of Yap1 in adult RGLs. Data information: Scale bars: 20 µm.

**Figure EV5.**
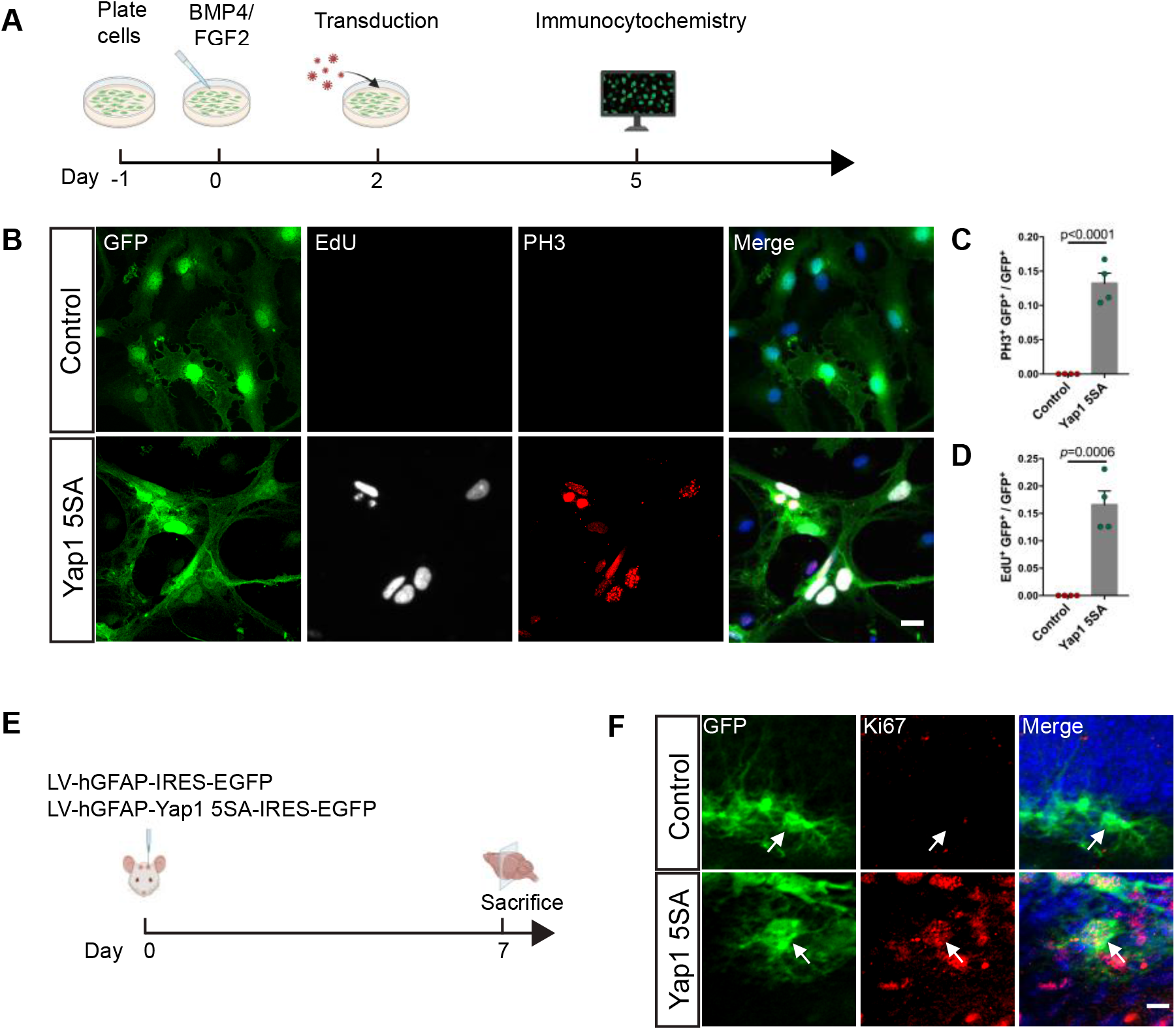
Overexpression of Yap1-5SA induces the proliferation of quiescent adult NSCs *in vitro* and astrocytes *in vivo*. A. Schematic diagram of the experimental design for the analysis of the effect of Yap1 overexpression in quiescent NSCs *in vitro*. B. Immunofluorescence for GFP, EdU (administered 6 hours before fixation) and PH3 (to identify proliferative cells) in control and Yap1-5SA overexpressing NSCs. C-D. Quantification of the data in (B). Yap1-5SA induces proliferation of quiescent NSCs *in vitro*. n=4 independent cultures for each group. E. Schematic diagram of the experimental design for the analysis of the effect of Yap1 overexpression in astrocytes in hilus. F. Immunofluorescence for GFP and Ki67 in coronal brain sections. Arrows indicate astrocytes in hilus. Data information: Data are represented as mean ± SEM. Unpaired Student’s t test. Scale bars: 20 µm.

**Figure EV6.**
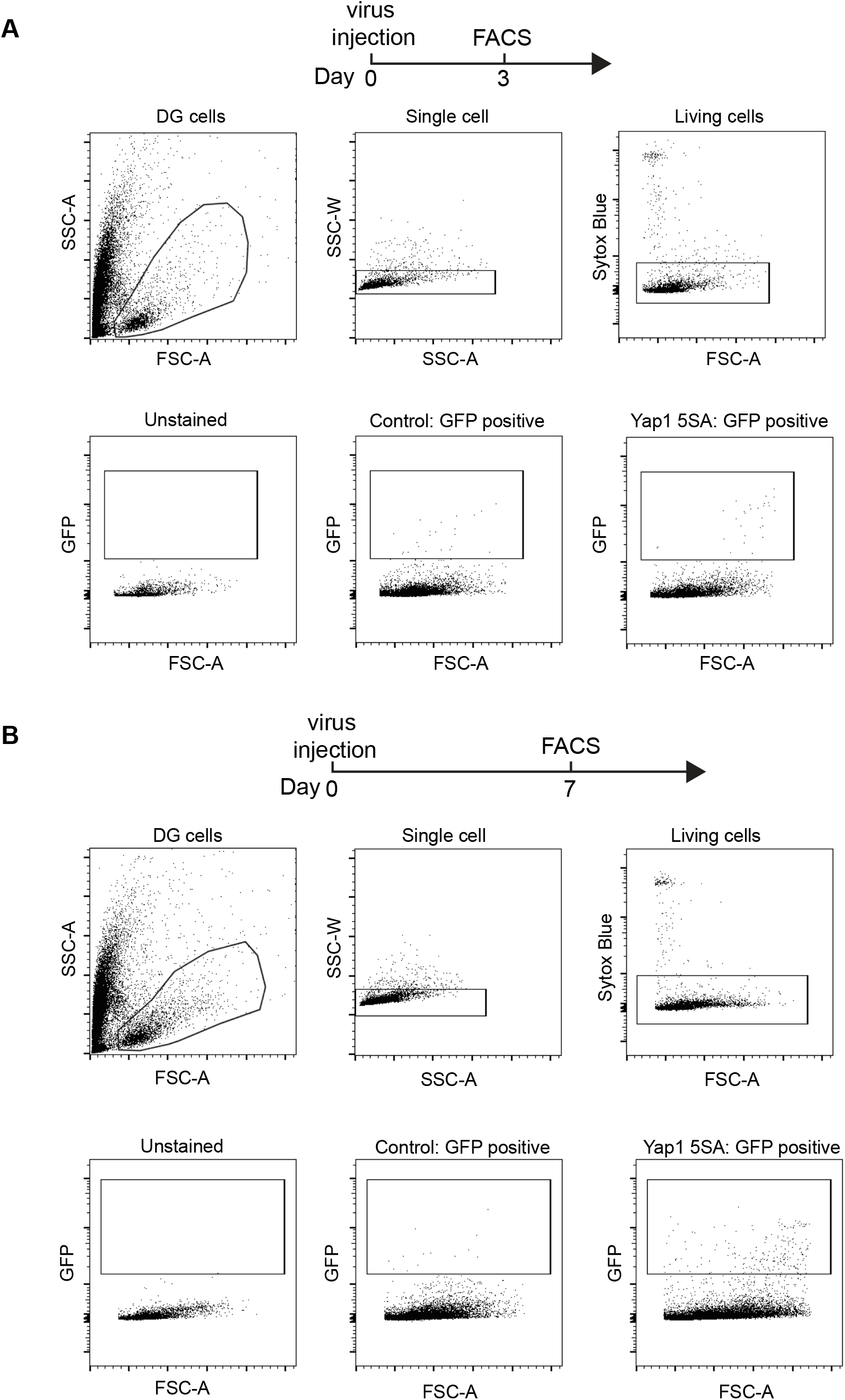
Gating strategy for FACS at 3 dpi and 7 dpi. A-B. FACS strategy to obtain GFP positive cells (transduced cells by LV-hGFAP-IRES-EGFP or LV-hGFAP-Yap1-5SA-IRES-EGFP). First gate uses stringent FSC/SSC gating to exclude cellular debris; second excludes cell aggregates to obtain single cells; third excludes dead and dying cells by Sytox Blue; fourth excludes GFP negative cells. GFP positive cells were collected in both control and Yap1 overexpression group at 3dpi and 7dpi.

**Figure EV7.**
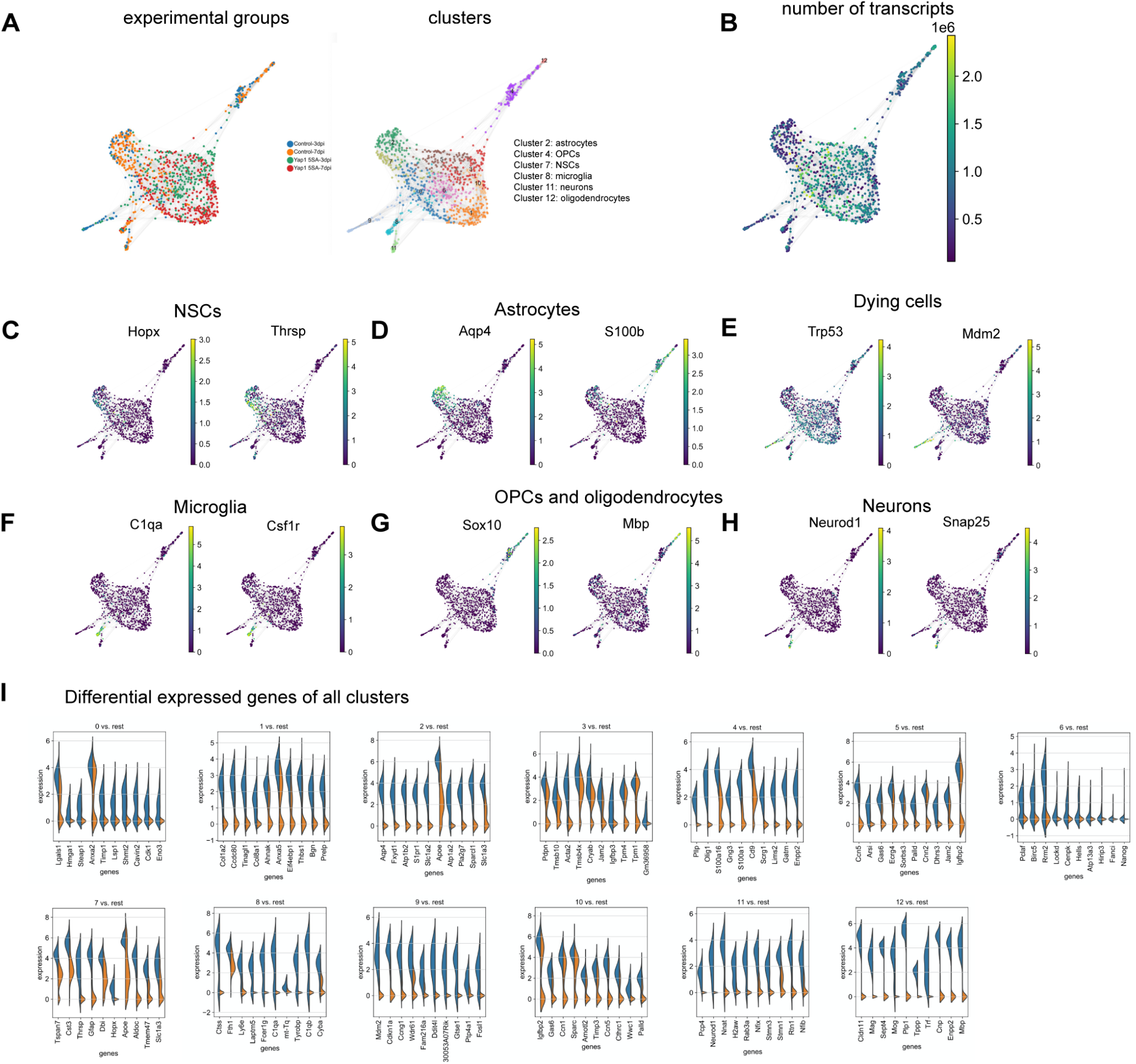
Clustering of single cell RNA sequencing data and cell type identification based on differential gene expression analysis. A. Experimental groups and 13 different cell clusters. B. Number of transcripts in both control and Yap1-5SA-expressing cells C-H. Markers for each cell type including NSCs, astrocytes, dying cells, microglia, oligodendroglial cells and neurons. I. Identification of top 10 markers for each cluster via differential gene expression analysis.

**Figure EV8.**
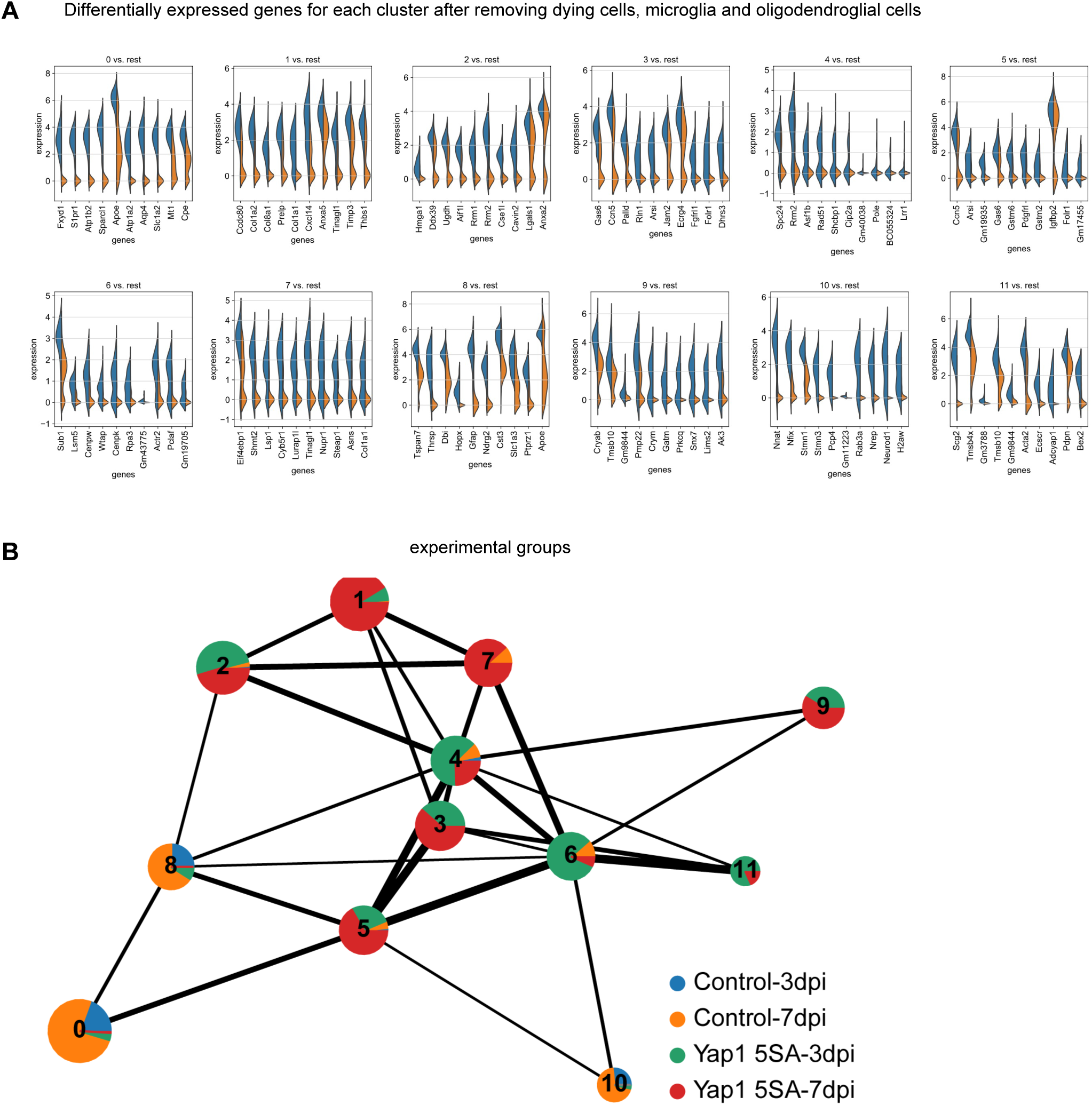
Re-clustering of the single cell RNA sequencing data after removal of dying cells, microglia and oligodendroglial cells. A. Top 10 markers for each cluster after re-clustering the data. B. Composition of experimental groups in each cluster are indicated in the plot.

## MATERIAL AND METHODS

### Animals and tamoxifen treatment

The study was performed in accordance with the guidelines of the German Animal Welfare Act and the European Directive 2010/63/EU for the protection of animals used for scientific purposes, and was approved by the Rhineland-Palatinate State Authority (permit number 23 177 07-G15-1-031). Mice were housed on a 12-h light/dark cycle in standard cages with free access to food and water. The study was conducted in both male and female mice. Wild type C57BL/6J mice were purchased from Janvier at 8 weeks of age (Janvier Labs). Yap1 conditional knockout mice (*GlastCre^ERT2^; Yap1 ^flox/flox^; CAG-CAT-EGFP*, referred to as Yap1 cKO) were generated by crossing the *GlastCre^ERT2^* with *Yap1 ^flox/flox^ (*Jackson laboratory Stock NO. 027929) and *CAG-CAT-EGFP* mouse lines.

Tamoxifen (Sigma, T5648) was dissolved in corn oil with 10 % ethanol to prepare a 20 mg/ml solution and administered once daily to both control (*GlastCre^ERT2^; Yap1^wt/wt^; CAG-CAT-EGFP*) and Yap1 cKO mice via intraperitoneal injections with 100 mg/kg for 5 consecutive days. Mice were sacrificed at indicated times as described in this article.

### Tissue preparation

Mice were deeply anesthetized with a combination of 120 mg/kg Ketamine (Zoetis) and 16 mg/kg Xylazine (Bayer) (in 0.9 % NaCl, i.p.) and transcardially perfused with saline, followed by 4 % paraformaldehyde (PFA, Sigma, P6148). Brains were harvested and post-fixed in 4 % PFA overnight at 4 °C. Coronal brain sections were prepared at a thickness of 50 µm using a Thermo Scientific vibrating blade microtome (Microm HM650V). Brain sections were stored in a cryoprotective solution (20 % glucose, 40 % ethylene glycol, 0.025 % sodium azide, and 0.05 M phosphate buffer, pH 7.4) at −20 °C.

### Plasmids cloning and lentivirus production

To generate lentiviral destination plasmids (LV-hGFAP-IRES-EGFP, LV-hGFAP-Yap1-IRES-EGFP, LV-hGFAP-Yap1(5SA)-IRES-EGFP), Yap1 and Yap1-5SA with attB sites were amplified PCR (polymerase chain reaction) from pQCXIH-Myc-Yap1(5SA) (a kind gift from Kunliang Guan, Addgene plasmid # 33093) and pcDNA3-Yap1 (a kind gift from Stefano Piccolo, University of Padova). The amplified fragments were cloned into pDNR221 to generate p-entry plasmids using Gateway cloning (ThermoFisher Scientific). Lentiviral destination plasmids were generated by the LR recombination between p-entry and destination plasmids.

Lentivirus production was performed as described previously (Tiscornia *et al.*, 2006). Second generation lentiviral packaging plasmids psPAX and VSV-G envelope expressing plasmid pMD2.G (kind gifts from Didier Trono, Addgene plasmid #12260 and #12259) were used for the virus production. Viral pellets were resuspended in phosphate-buffered saline (PBS) and stored at −80°C.

### Lentivirus injection

Stereotactic injections of lentivirus were performed in 8 week-old C57BL/6J mice. Prior to stereotactic injections and on the day following the surgery, mice were treated with analgesics (Rimadyl®, Zoetis, 4 mg/Kg of body weight in 0.9 % NaCl) by intraperitoneal injection and then anesthetized with a combination of 0.5 mg/kg Medetomidin (Pfizer), 5mg/kg Midazolam (Hameln) and 0.025 mg/kg Fentanyl (Albrecht) (in 0.9 % NaCl). Mice were placed in a stereotactic frame and kept on an animal heating pad to control body temperature during surgery. A small craniotomy was performed and 1 µl of the lentivirus was slowly injected using a pulled glass capillary (Hirschmann, 9600105) into the dentate gyrus at the following coordinates relative to Bregma: −2.0 AP (anterior-posterior), 1.4 ML (medio-lateral), −2.0 DV (dorso-ventral). The capillary was left in place for another 5 minutes before withdrawal to allow diffusion of the virus. Then anesthesia was antagonized by intraperitoneal injection of 2.5 mg/kg Atipamezol (Pfizer), 0.5 mg/kg Flumazenil (Hameln) and 0.1 mg/kg Buprenorphin (RB Pharmaceutials) (in 0.9 % NaCl). Animals were allowed to recover, returned to their home cages, and their physical condition was monitored daily.

### Adherent adult neural stem cell culture

Adult neural stem cell cultures were prepared from the DG of young adult (8 week-old) mice as previously reported with minor modifications (Babu *et al.*, 2011). Briefly, DG was dissected from adult mouse brain in calcium and magnesium-free Hank's Balanced Salt Solution (HBSS, ThermoFisher Scientific, cat# 14170112) and the tissue was enzymatically dissociated using the Neural Tissue Dissociation Kit (Miltenyi Biotec, Germany, cat# 130-092-628) according to the manufacturer instructions. For the Fluorescence Activated Cell Sorting (FACS), the cell pellet was resuspended with PBS containing 5 % Fetal bovine serum (FBS, Life Technology, cat# 10270-106).

For the cell culture, the cells were purified as previously described (Ortega et al., 2013). The cell pellet was finally resuspended in Neurobasal Glutamax (Invitrogen, cat# 21103049) supplemented with B27 (Invitrogen, cat# 17504001), 20 ng/ml epidermal growth factor (EGF, Peprotech, cat#315-09), 20 ng/ml fibroblast growth factors (FGF2, Peprotech, cat# 100-18B), 100 units/ml penicillin and 100 μg/ml streptomycin (Invitrogen, cat# 15140122), and cells were plated in Poly-D-lysine hydrobromide (PDL, Sigma, cat# P0899) pre-coated 24-well plates (VWR, cat# 7340020). For the NSC passaging, the cells were trypsinized using 0.05 % Trypsin-EDTA (ThermoFisher Scientific, cat# 25300054) for 5 minutes at 37°C. Neural stem cells were used for experiments for up to 10 passages.

### Immunohistochemistry and immunocytochemistry

For immunohistochemistry, brain sections were blocked with 5 % donkey serum (Sigma, S30) and 0.5 % Triton-100 (Sigma, cat# X100-500ml) in Tris-buffered saline (TBS, 50 mM Tris-Cl, pH 7.5; 150 mM NaCl) and stained with primary antibodies overnight at 4°C followed by secondary antibodies for 1 h at room temperature. Primary antibodies used in this study were as follows: goat anti-DCX (1:300, Santa Cruz, sc-8066), guinea-pig anti-DCX (1:500, Millipore, AB2253), mouse anti-GFAP (1:400, Millipore, MAB360), goat anti-GFAP (1:800, Abcam, ab53554), chicken anti-GFP (1:1000, Aves Lab, GFP-1020), rabbit anti-Mcm2 (1:800, Cell Signaling, 4007S), rabbit anti-Olig2 (1:1500, Millipore, AB9610), goat anti-Sox2 (1:500, Santa Cruz, sc-17320), rabbit anti-Sox2 (1:500, Abcam, ab137385), rabbit anti-Yap1 (1:200, Cell Signaling, 14074S), mouse anti-Yap1 (1:200, Sigma, WH0010413M1). Secondary antibodies used were as follows: donkey anti-chicken Alexa 488 (1:200, Jackson ImmunoResearch, 703545155), donkey anti-goat Cy3 (1:500, Dianova, 705165147), donkey anti-goat Cy5 (1:300, Dianova, 705165147), donkey anti-Guinea pig Cy5 (1:300, Dianova, 705175148), donkey anti-mouse Alexa 647 (1:300, Invitrogen, A31571), donkey anti-rabbit Cy3 (1:500, Dianova, 711165152).

Images were acquired with a LEICA TCS SPE confocal microscope equipped with a 40x oil objective (NA 1.3) or an upright ZEISS Axio Imager.M2 epifluorescent microscope equipped with a 40x dry objective (NA 0.75) and an ApoTome.2 system (Zeiss GmbH). 4 −5 images were taken from every brain for quantifications. Images were analyzed using Image J software (NIH).

For immunocytochemistry, adult neural stem cells were fixed in 4 % PFA for 10 mins at room temperature, followed by washing with PBS for 3 times. Cells were blocked with 2 % bovine serum albumin (BSA) (Sigma, A9418) and 0.2 % Triton-100 for 45 min in TBS and incubated with primary antibodies overnight at 4 °C followed by incubation with secondary antibodies for 1 h at room temperature. For BrdU (5-bromo-2'-deoxyuridine, Sigma, cat# B5002) staining, cells were incubated in 2 N HCl at 37°C for 10 min, followed by 0.1 M boric acid for 10 min at room temperature. EdU (5-ethynyl-2’-deoxyuridine, ThermoFisher Scientific, cat# A10044) staining was performed by Click-it Alexa Fluor 647 imaging kit (Thermo Fisher Scientific, cat# C10086) according to the manufacturer instructions. Primary antibodies used in this study were as follows: rat anti-BrdU (1:500, ABD Serotec, OBT0030), chicken anti-GFP (1:1000, Aves Lab, GFP-1020), rabbit anti-Nestin (1:500, BioLegend, 839801), rabbit anti-PH3 (1:300, Millipore, 06570), rabbit anti-Yap1 (1:200, Cell Signaling, 14074S), mouse anti-Yap1 (1:200, Sigma, WH0010413M1). Secondary antibodies used were as follows: donkey anti-chicken Alexa 488 (1:1000, Jackson ImmunoResearch, 703545155), donkey anti-rabbit Cy3 (1:1000, Dianova, 711165152), donkey anti-rat Alexa 547 (1:1000, Interchim, FP-SB6120).

Images of immunostained cells in culture were acquired with an upright ZEISS Axio Imager.M2 epifluorescent microscope equipped with 40x dry objective (NA 0.75) and an ApoTome.2 system (Zeiss GmbH). 4-5 images were taken from each independent experiment for the quantifications. Images were analyzed using Image J software (NIH).

### Flow cytometry

For flow cytometry of lentivirus transduced cells, animals were sacrificed by cervical dislocation and their brains immediately placed on ice. The DG was microdissected and the tissue was enzymatically dissociated using the Neural Tissue Dissociation Kit (Miltenyi Biotec, Germany, cat# 130-092-628) according to the manufacturer instructions. Sytox Blue (1/1000, Life Technologies, cat# S34857) was added to the cell suspension prior to sorting to exclude dead cells. Gates and compensations were set before every experiment using control samples without any fluorochromes. Single cell suspensions were sorted in a FACSAria II SORP (BD Bioscience) using a 100 μm nozzle at 20 psi. EGFP-positive cells from control and Yap1-5SA groups were directly collected into ice-cold lysis buffer (0.2 %Triton X-100, containing RNAse inhibitors, Clontech, cat# 635013) in 96-well plates (Eppendorf, cat# 0030128680) and frozen at −80°C until library preparation.

### Library preparation and single cell RNA sequencing

An adapted protocol from the Smart-seq2 method (Picelli *et al.*, 2014) was used. Upon thawing the plates on ice, reverse transcription was performed using an oligo(dT) primer and a locked nucleic acid (LNA)-containing template-switching oligonucleotide (IDT; custom oligo). For all samples, we included ERCC spike-in controls (Ambion, cat#4456740) at a 1:10^6^ dilution. Full-length cDNAs were amplified by 18 cycles of PCR using KAPA HiFi DNA polymerase (Roche, cat#7958935001). Amplified cDNAs for a set of random selected samples were quantified and run on a High Sensitivity Bioanalyzer chip (Agilent, cat# 5067-4626) on a 2100 Bioanalyzer (Agilent).

Libraries were prepared using the Illumina Nextera XT DNA sample preparation kit (Illumina, cat#FC-131-1096) with a minor modification. The modification is based on the Tagmentation protocol from Fluidigm, which uses 25 % of the reagents per tagmentation reaction with a starting amount of 100-125 pg of cDNA. Libraries were amplified in 12 PCR cycles. Three microliters of each library were pooled into a single tube and the pool was subsequently bead-purified using Agencout AMPure XP beads (Beckman Coulter GmbH, cat# A63882). The purified pool was profiled in a High Sensitivity DNA chip on a 2100 Bioanalyzer (Agilent) and quantified using the Qubit dsDNA HS Assay Kit (Invitrogen, cat#Q32854) in a Qubit 2.0 Fluorometer (Life technologies). Four pools (each one from one of four 96-well plates) were pooled together at equal concentrations and sequenced on a single NextSeq 500 High Output Flow Cell with single reads for 1 × 75 cycles for read 1 plus 2 × 8 cycles for each index read.

### Single cell RNA sequencing data processing

All downstream analysis was performed using the open-source R software accessed via RStudio server (R version 3.6.3). Smart-Seq2 raw data demultiplexing was performed using Illumina’s bcl2fastq conversion software v.2.19.1 and overall sequence quality was assessed with FastQC v0.11.5. STAR v.2.5.2b with default parameters (except –outFilterMismatchNoverLmax 0.04) was used to align reads to the mouse reference genome GRCm38 (mm10), supplemented with ERCC RNA Spike-In Mix (ThermoFischer) control sequences and considering the GTF gene annotation from Ensembl release 88. Read summarization at the gene level was performed using Subread featureCounts v.1.6.2 with default parameters.

The quality control and normalization pipeline were applied as previously described (Amezquita *et al.*, 2020). Briefly, cells with log transformed library sizes and number of genes below 3 median absolute deviations (MADs) from the median of the respective distributions were removed. In addition, we removed cells in which the proportion of mitochondrial genes and spike-in RNAs was 3 MADs above the respective median proportions. Lastly, genes with average counts across samples below a threshold of 1 were filtered out.

### Single cell transcriptome analysis

Using python (3.8.1) and standard python libraries (ipykernel 5.3.0. matplotlib 3.2.2, numpy 1.17.3, pandas 1.0.5, scipy 1.4.1), we employed scanpy (1.5.1), anndata (0.7.3) and anndata2ri (1.0.2) (Wolf *et al.*, 2018) (Wolf et al., 2018) to perform single cell analysis. We filtered out cells with less than 500 or more than 6000 genes detected and cells with less than 5 % mitochondrial genes. Moreover, we filtered out genes that are expressed in less than 10 cells. After normalization and log transformation we regressed out unwanted sources of variation from differences in cell cycle scores, percent mitochondrial genes, number of transcripts per cell and percentage of spike-in. We identified highly variable genes using default parameters which were further used for exploratory analysis via principal component analysis and force-directed graph embedding (python-igraph 0.8.2). Clustering of cells was done using the leiden-algorithm (0.8.1) implementation within scanpy. Differential gene expression analysis between Leiden clusters was performed using function sc.tl.rank_genes_groups with wilcoxon testing. For quantification of RNA splicing from the raw sequencing data we used velocyto (0.17) (La Manno *et al.*, 2018) with default smartseq2 parameters. Subsequently we employed scvelo (0.2.3) (Bergen *et al.*, 2020) for RNA velocity estimations using the stochastic algorithm with default parameters after recovering the dynamics and using latent time. For the estimation of a trajectory pseudotime we employed the diffusion pseudotime algorithm implementation in scvelo.

The analysis of adult hippocampal dataset (Hochgerner *et al.*, 2018) was performed using the open-source R software accessed via RStudio server (R version 3.6.3). Briefly, normalization was performed using SCTransform with default parameters in Seurat (v3.1). We selected different cut-offs of the number of PCs and empirically found that downstream clustering analyses were optimized when using a 15-PC cutoff. The first 15 PCs were selected and used for two-dimension Uniform Manifold Approximation and Projection (UMAP), implemented by the Seurat software with default parameters. Based on the UMAP map, 5 clusters were identified using the function FindClusters in Seurat with the resolution parameter of 0.5. Radial glia-like cells, astrocytes, nIPCs and neuroblasts were identified using FindAllMakers with default parameters. Differentially expressed genes that were expressed in at least 25 % cells within the cluster and with a fold change more than 0.5 (log scale) were considered to be marker genes. To calculate Yap signature and active signature, AddModuleScore function was used in Seurat with default parameters.

### Statistics

Statistical analysis was performed by two-tailed unpaired Student’s t test using GraphPad Prism version 7 (GraphPad Software, Inc.). Data are represented as mean ± SEM. P values are indicated in the graphs.

## Data availability

The adult hippocampal dataset is available at GEO accession GSE95753. Sequencing data have been deposited at GEO accession GSE144967.

## Acknowledgements

This work was supported by grants from the German Research Foundation (CRC1080, project number 221828878, CRC1193 project number 26410226), Wellcome Trust (206410/Z/17/Z) to BB; core funding to the Francis Crick Institute from Cancer Research UK, The Medical Research Council and the Wellcome Trust (FC001002); funding from the Naturwissenschaftlich-Medizinischen Forschungszentrum (NMFZ) of the Johannes Gutenberg University Mainz to CB and BB. J.J-A was supported by research fellowships from Fundación Alfonso Martín Escudero, Fundación Ramón Areces and the European Commission (Marie Skłodowska-Curie Action).

We are grateful to Flow Cytometry Core Facility, Genomic Core Facility, Microscopy Core Facility and Bioinformatics Core Facility of the Institute of Molecular Biology (IMB) Mainz for their support regarding single cell RNA sequencing experiments and raw data processing. Parts of schematic diagram were created with BioRender.com.

## Author contributions

Most of the data were generated and analyzed by WF; JJ-A contributed to in vivo experiments and analysis; WF, GA-L, MA-N and SF performed bioinformatics analysis; Experimental design: BB, CB, SP, WF; Project conceptualization and oversight: BB; Manuscript writing: BB, JJ-A, WF; all authors read and approved the final manuscript.

## References

Amezquita, RA, Lun, AT, Becht, E, Carey, VJ, Carpp, LN, Geistlinger, L, Marini, F, Rue-Albrecht, K, Risso, D, Soneson, C (2020) Orchestrating single-cell analysis with Bioconductor. Nature methods 17: 137–145

Andersen, J, Urbán N, Achimastou, A, Ito, A, Simic, M, Ullom, K, Martynoga, B, Lebel, M, Göritz, C, Frisén J (2014) A transcriptional mechanism integrating inputs from extracellular signals to activate hippocampal stem cells. Neuron 83: 1085–1097

Babu, H, Claasen J-H, Kannan, S, Rünker, AE, Palmer, T, Kempermann, G (2011) A protocol for isolation and enriched monolayer cultivation of neural precursor cells from mouse dentate gyrus. Frontiers in neuroscience 5:89

Berg, DA, Su, Y, Jimenez-Cyrus, D, Patel, A, Huang, N, Morizet, D, Lee, S, Shah, R, Ringeling, FR, Jain, R (2019) A common embryonic origin of stem cells drives developmental and adult neurogenesis. Cell 177: 654–668. e15

Bergen, V, Lange, M, Peidli, S, Wolf, FA, Theis, FJ (2020) Generalizing RNA velocity to transient cell states through dynamical modeling. Nature biotechnology 38: 1408–1414

Cappello, S, Gray, MJ, Badouel, C, Lange, S, Einsiedler, M, Srour, M, Chitayat, D, Hamdan, FF, Jenkins, ZA, Morgan, T (2013) Mutations in genes encoding the cadherin receptor-ligand pair DCHS1 and FAT4 disrupt cerebral cortical development. Nature genetics 45: 1300–1308

Castellan, M, Guarnieri, A, Fujimura, A, Zanconato, F, Battilana, G, Panciera, T, Sladitschek, HL, Contessotto, P, Citron, A, Grilli, A (2021) Single-cell analyses reveal YAP/TAZ as regulators of stemness and cell plasticity in glioblastoma. Nature cancer 2: 174–188

Cordenonsi, M, Zanconato, F, Azzolin, L, Forcato, M, Rosato, A, Frasson, C, Inui, M, Montagner, M, Parenti, AR, Poletti, A (2011) The Hippo transducer TAZ confers cancer stem cell-related traits on breast cancer cells. Cell 147: 759–772

Ding, R, Weynans, K, Bossing, T, Barros, CS, Berger, C (2016) The Hippo signalling pathway maintains quiescence in Drosophila neural stem cells. Nature communications 7: 1–12

Dulken, BW, Buckley, MT, Negredo, PN, Saligrama, N, Cayrol, R, Leeman, DS, George, BM, Boutet, SC, Hebestreit, K, Pluvinage, JV (2019) Single-cell analysis reveals T cell infiltration in old neurogenic niches. Nature 571: 205–210

Engler, A, Rolando, C, Giachino, C, Saotome, I, Erni, A, Brien, C, Zhang, R, Zimber-Strobl, U, Radtke, F, Artavanis-Tsakonas S (2018) Notch2 signaling maintains NSC quiescence in the murine ventricular-subventricular zone. Cell reports 22: 992–1002

Gil-Ranedo, J, Gonzaga, E, Jaworek, KJ, Berger, C, Bossing, T, Barros, CS (2019) STRIPAK members orchestrate hippo and insulin receptor signaling to promote neural stem cell reactivation. Cell reports 27: 2921–2933. e5

Hamon, A, García-García, D, Ail, D, Bitard, J, Chesneau, A, Dalkara, D, Locker, M, Roger, JE, Perron, M (2019) Linking YAP to Müller glia quiescence exit in the degenerative retina. Cell reports 27: 1712–1725. e6

Han, J, Calvo, C-F, Kang, TH, Baker, KL, Park, J-H, Parras, C, Levittas, M, Birba, U, Pibouin-Fragner, L, Fragner, P (2015) Vascular endothelial growth factor receptor 3 controls neural stem cell activation in mice and humans. Cell reports 10: 10–1158

Hao, Y, Hao, S, Andersen-Nissen, E, Mauck III, WM, Zheng, S, Butler, A, Lee, MJ, Wilk, AJ, Darby, C, Zager, M (2021) Integrated analysis of multimodal single-cell data. Cell

Harada, Y, Yamada, M, Imayoshi, I, Kageyama, R, Suzuki, Y, Kuniya, T, Furutachi, S, Kawaguchi, D, Gotoh, Y (2021) Cell cycle arrest determines adult neural stem cell ontogeny by an embryonic Notch-nonoscillatory Hey1 module. Nature communications 12: 1–16

Harris, L, Rigo, P, Stiehl, T, Gaber, ZB, Austin, SH, del, Mar Masdeu, M, Edwards, A, Urbán, N, Marciniak-Czochra, A, Guillemot, F (2021) Coordinated changes in cellular behavior ensure the lifelong maintenance of the hippocampal stem cell population. Cell stem cell 28: 863–876. e6

Hochgerner, H, Zeisel, A, Lönnerberg, P, Linnarsson, S (2018) Conserved properties of dentate gyrus neurogenesis across postnatal development revealed by single-cell RNA sequencing. Nature neuroscience 21: 290–299

Ibrayeva, A, Bay, M, Pu, E, Jörg, DJ, Peng, L, Jun, H, Zhang, N, Aaron, D, Lin, C, Resler, G (2021) Early stem cell aging in the mature brain. Cell stem cell 28: 955–966. e7

Kalamakis, G, Brüne, D, Ravichandran, S, Bolz, J, Fan, W, Ziebell, F, Stiehl, T, Catalá-Martinez, F, Kupke, J, Zhao, S (2019) Quiescence modulates stem cell maintenance and regenerative capacity in the aging brain. Cell 176: 1407–1419. e14

Kawai, H, Kawaguchi, D, Kuebrich, BD, Kitamoto, T, Yamaguchi, M, Gotoh, Y, Furutachi, S (2017) Area-specific regulation of quiescent neural stem cells by Notch3 in the adult mouse subependymal zone. Journal of Neuroscience 37: 11867–11880

Kostic, M, Paridaen, JT, Long, KR, Kalebic, N, Langen, B, Grübling, N, Wimberger, P, Kawasaki, H, Namba, T, Huttner, WB (2019) YAP activity is necessary and sufficient for basal progenitor abundance and proliferation in the developing neocortex. Cell reports 27: 1103–1118. e6

La Manno, G, Soldatov, R, Zeisel, A, Braun, E, Hochgerner, H, Petukhov, V, Lidschreiber, K, Kastriti, ME, Lönnerberg, P, Furlan, A (2018) RNA velocity of single cells. Nature 560: 494–498

Lavado, A, Park, JY, Paré J, Finkelstein, D, Pan, H, Xu, B, Fan, Y, Kumar, RP, Neale, G, Kwak, YD (2018) The Hippo pathway prevents YAP/TAZ-driven hypertranscription and controls neural progenitor number. Developmental cell 47: 576–591. e8

Lie D-C, Colamarino, SA, Song, H-J, Désiré L, Mira, H, Consiglio, A, Lein, ES, Jessberger, S, Lansford, H, Dearie, AR (2005) Wnt signalling regulates adult hippocampal neurogenesis. Nature 437: 1370–1375

Martynoga, B, Mateo, JL, Zhou, B, Andersen, J, Achimastou, A, Urbán, N, van, den Berg, D, Georgopoulou, D, Hadjur, S, Wittbrodt, J (2013) Epigenomic enhancer annotation reveals a key role for NFIX in neural stem cell quiescence. Genes & development 27: 1769–1786

Mira, H, Andreu, Z, Suh, H, Lie, DC, Jessberger, S, Consiglio, A, San Emeterio, J, Hortigüela, R, Marqués-Torrejón MÁ, Nakashima, K (2010) Signaling through BMPR-IA regulates quiescence and long-term activity of neural stem cells in the adult hippocampus. Cell stem cell 7: 78–89

Mo, JS, Park, HW, Guan, KL (2014) The Hippo signaling pathway in stem cell biology and cancer. EMBO reports 15: 642–656

Mori, T, Tanaka, K, Buffo, A, Wurst, W, Kühn, R, Götz M (2006) Inducible gene deletion in astroglia and radial glia—a valuable tool for functional and lineage analysis. Glia 54: 21–34

Moya, IM, Halder, G (2019) Hippo–YAP/TAZ signalling in organ regeneration and regenerative medicine. Nature Reviews Molecular Cell Biology 20: 211–226

Mukhtar, T, Breda, J, Grison, A, Karimaddini, Z, Grobecker, P, Iber, D, Beisel, C, van Nimwegen, E, Taylor, V (2020) Tead transcription factors differentially regulate cortical development. Scientific reports 10: 1–19

Nakamura, T, Colbert, MC, Robbins, J (2006) Neural crest cells retain multipotential characteristics in the developing valves and label the cardiac conduction system. Circulation research 98: 1547–1554

Picelli, S, Faridani, OR, Björklund ÅK, Winberg, G, Sagasser, S, Sandberg, R (2014) Full-length RNA-seq from single cells using Smart-seq2. Nature protocols 9: 171–181

Plouffe, SW, Lin, KC, Moore, JL, Tan, FE, Ma, S, Ye, Z, Qiu, Y, Ren, B, Guan, K-L (2018) The Hippo pathway effector proteins YAP and TAZ have both distinct and overlapping functions in the cell. Journal of Biological Chemistry 293: 11230–11240

Rueda, EM, Hall, BM, Hill, MC, Swinton, PG, Tong, X, Martin, JF, Poché RA (2019) The Hippo pathway blocks mammalian retinal Müller glial cell reprogramming. Cell reports 27: 1637–1649. e6

Shin, J, Berg, DA, Zhu, Y, Shin, JY, Song, J, Bonaguidi, MA, Enikolopov, G, Nauen, DW, Christian, KM, Ming, G-l (2015) Single-cell RNA-seq with waterfall reveals molecular cascades underlying adult neurogenesis. Cell stem cell 17: 360–372

Song, J, Zhong, C, Bonaguidi, MA, Sun, GJ, Hsu, D, Gu, Y, Meletis, K, Huang, ZJ, Ge, S, Enikolopov, G (2012) Neuronal circuitry mechanism regulating adult quiescent neural stem-cell fate decision. Nature 489: 150–154

Tiscornia, G, Singer, O, Verma, IM (2006) Production and purification of lentiviral vectors. Nature protocols 1: 241–245

Urbán, N, Blomfield, IM, Guillemot, F (2019) Quiescence of adult mammalian neural stem cells: a highly regulated rest. Neuron 104: 834–848

Urbán, N, van, den Berg, DL, Forget, A, Andersen, J, Demmers, JA, Hunt, C, Ayrault, O, Guillemot, F (2016) Return to quiescence of mouse neural stem cells by degradation of a proactivation protein. Science 353: 292–295

Wexler, EM, Paucer, A, Kornblum, HI, Palmer, TD, Geschwind, DH (2009) Endogenous Wnt signaling maintains neural progenitor cell potency. Stem cells 27: 1130–1141

Wolf, FA, Angerer, P, Theis, FJ (2018) SCANPY: large-scale single-cell gene expression data analysis. Genome biology 19: 1–5

Yu F-X, Guan, K-L (2013) The Hippo pathway: regulators and regulations. Genes & development 27: 355–371

Zhang, N, Bai, H, David, KK, Dong, J, Zheng, Y, Cai, J, Giovannini, M, Liu, P, Anders, RA, Pan, D (2010) The Merlin/NF2 tumor suppressor functions through the YAP oncoprotein to regulate tissue homeostasis in mammals. Developmental cell 19: 27–38

Zhang, R, Boareto, M, Engler, A, Louvi, A, Giachino, C, Iber, D, Taylor, V (2019) Id4 downstream of Notch2 maintains neural stem cell quiescence in the adult hippocampus. Cell reports 28: 1485–1498. e6

Zhao, B, Wei, X, Li, W, Udan, RS, Yang, Q, Kim, J, Xie, J, Ikenoue, T, Yu, J, Li, L (2007) Inactivation of YAP oncoprotein by the Hippo pathway is involved in cell contact inhibition and tissue growth control. Genes & development 21: 2747–2761

